# Electron Cryo-Tomography Reveals the *Caulobacter vibrioides* Tight Adherence Pilus Architecture

**DOI:** 10.1101/2025.11.06.686991

**Authors:** Stefano Maggi, Rachel Rosenzweig, Gaël Panis, Patrick H. Viollier, Grant J. Jensen

**Affiliations:** Department of Chemistry and Biochemistry, Brigham Young University, Provo, UT 84604, USA; Division of Biology, California Institute of Technology, Pasadena, CA, USA; Department of Microbiology and Molecular Medicine, University of Geneva, Geneva, Switzerland

**Keywords:** Cryo-electron tomography, cryo-ET, *Caulobacter crescentus*, Tight adherence pilus, Tad pilus, Type IV pilus, T4P, Type II secretion system, T2SS

## Abstract

The type IV filament superfamily is a widespread group of molecular machines involved in natural transformation, motility, adhesion, nutrient uptake, and secretion of a wide spectrum of protein substrates. The gram-negative bacterium *Caulobacter vibrioides* expresses the tight adherence (Tad) pilus, a type IV machine involved in surface colonization. Here we investigated the proteins involved in Tad pilus production by ΦCbK resistance screening and the order of machine assembly and its polar remnant by fluorescence tagging. Using electron cryo-tomography and subtomogram averaging of wild-type and mutant strains, we resolved the Tad pilus machine architecture and built an integrative model of its structure. The resulting model suggests the individual roles of multiple Tad proteins. Together the data also reveals the Tad pilus machine’s assembly order.

## INTRODUCTION

Pili, the hair-like appendages found on the surface of Archaea and Bacteria, fall into several filament superfamilies. Type IV pili are involved in multiple cellular functions such as surface adhesion (Piepenbrink and Sundberg 2016), motility (Burrows 2012), genetic exchange (Meibom et al. 2005), and biofilm formation (Pohlschroder and Esquivel 2015). A unique member of the type IV pilus family is the tight-adherence (Tad) pilus, which originated in Archaea and was later acquired by bacteria (Denise et al. 2019).

In *Caulobacter vibrioides* the Tad pilus plays a key role in surface adhesion and has a complex spatiotemporal regulation that links surface attachment with cell cycle differentiation (Mignolet et al. 2018; Del Medico et al. 2020). The *C. vibrioides* cell cycle is characterized by an alternation of two asymmetric cell types: stalked and swarmer (Collier 2019; Schrader et al. 2016; Skerker and Laub 2004). Swarmer cells, which possess both a polar flagellum and Tad pili, swim until they encounter a nutrient-rich surface. The Tad pili mediate initial attachment, draw the cell close, and orient it upright (Sangermani et al. 2019). In this configuration the C-di-GMP levels increase (Sangermani et al. 2019; Snyder et al. 2020), activating a profound remodeling in which both the flagellum and Tad pili are dismantled and a polysaccharide-based adhesin (holdfast) is subsequently secreted, permanently anchoring the cell to the surface. This pole undergoes further modifications resulting in the formation of the stalk, a cylindrical extension of the cell envelope, whose exact function is still unclear (Barrows and Goley 2023; Klein et al. 2013).

At least three main proteins regulate protein localization at the poles to ensure temporal and spatial control: PopZ–CckA (Holmes et al. 2016), SpmX–DivJ (Perez et al. 2017), and PodJ–PleC (Hinz et al. 2003). The latter is directly involved in Tad polar localization (Viollier et al. 2002). The PodJ–PleC signal is further reinforced by a second pathway involving ZitP and CpaM (Mignolet et al. 2016).

The bacteriophage ΦCbK requires Tad pili to infect *C. vibriodes*. Screening for ΦCbK resistance phenotype therefore identified the genes involved in Tad assembly (Skerker and Shapiro, n.d.; Christen et al. 2016). Thirteen of these genes are organized in a single operon: *cpaA*–*cpaK, cpaO*, and *pilA*; two others are distal: *cpaL* and *cpaM*.

Based on homology to other type IV pilus systems, hypothetical models of the Tad machinery have been proposed (Mignolet et al. 2018; Sangermani et al. 2019; Tomich et al. 2007; Whitfield and Brun 2024). These models suggest a hierarchical assembly starting with PodJ and ZitP recruiting CpaE and CpaM, respectively. CpaE is required for CpaC (secretin) assembly at the pole (Viollier 2002).

The structures of the *Pseudomonas aeruginosa* CpaC-CpaO complex and CpaB ring have been solved by cryo-EM (Tassinari et al. 2023; Evans et al. 2024). CpaO acts as CpaC’s pilotin while CpaB could function as an alignment complex. CpaF, the best-characterized Tad component, is the pilus fiber extension and retraction ATPase (Ellison et al. 2019; Hohl et al. 2024). The pilus structure has been recently solved and consists of a helical fiber of PilA monomers (Sonani et al. 2023). CpaI is present mainly in Alphaproteobacteria (Whitfield and Brun 2024) and is involved in Tad assembly (Christen et al. 2016), but its exact function remains unclear.

Here we investigated the proteins involved in Tad machine assembly and function by ΦCbK resistance assays, fluorescence microscopy, and electron cryo-tomography (cryo-ET) using new in-frame deletion mutants. Using these results and taking advantage of recent advances in protein structure prediction, we built an integrative model of the Tad machine. Our results identify which proteins form the Tad pilus machine, allowing us to solve its molecular architecture, identify its possible phage receptor, and provide support for a mixed-direction assembly mechanism.

## RESULTS

### Predicted localization of Tad proteins

For each of the fifteen Tad pilus machine proteins, we inferred its localization, domain architecture (Fig. S1), and interactions with other proteins based on sequence analysis and homology to previously characterized type IV pilus systems. We then built a new preliminary schematic model of the Tad pilus machine (Fig. 1). Our results suggest the following: CpaB has an inner transmembrane helix at its N-terminus. CpaD has a strong lipidation site that likely anchors the protein to the outer membrane, much like other type IV secretin-associated proteins (McCallum et al. 2021). CpaI has a periplasmic secretion signal peptide (SSP) and shares the same domain with the N-terminus of CpaC (Fig. S1, T2SS-T3SS_pil_N, IPR032789). We positioned the minor pilins CpaJ, CpaK, and CpaL at the tip of the pilus, as seen in other type IV pilus systems (Chang et al. 2016; Jacobsen et al. 2020). Homologs of CpaJ and CpaK in *Actinobacillus actinomycetemcomitans* both have a pilin SSP recognized by a prepilin peptidase and are directly involved in pilus biogenesis (Tomich et al. 2006), which further supports this positioning. While we wrote this manuscript, a review by Whitfield and Brun (2024) inferred these same locations of CpaB, CpaD, CpaI, CpaJ, CpaK, and CpaL. Our bioinformatics-based Tad pilus machine protein localization predictions agree with their new model.

**Fig 1.**
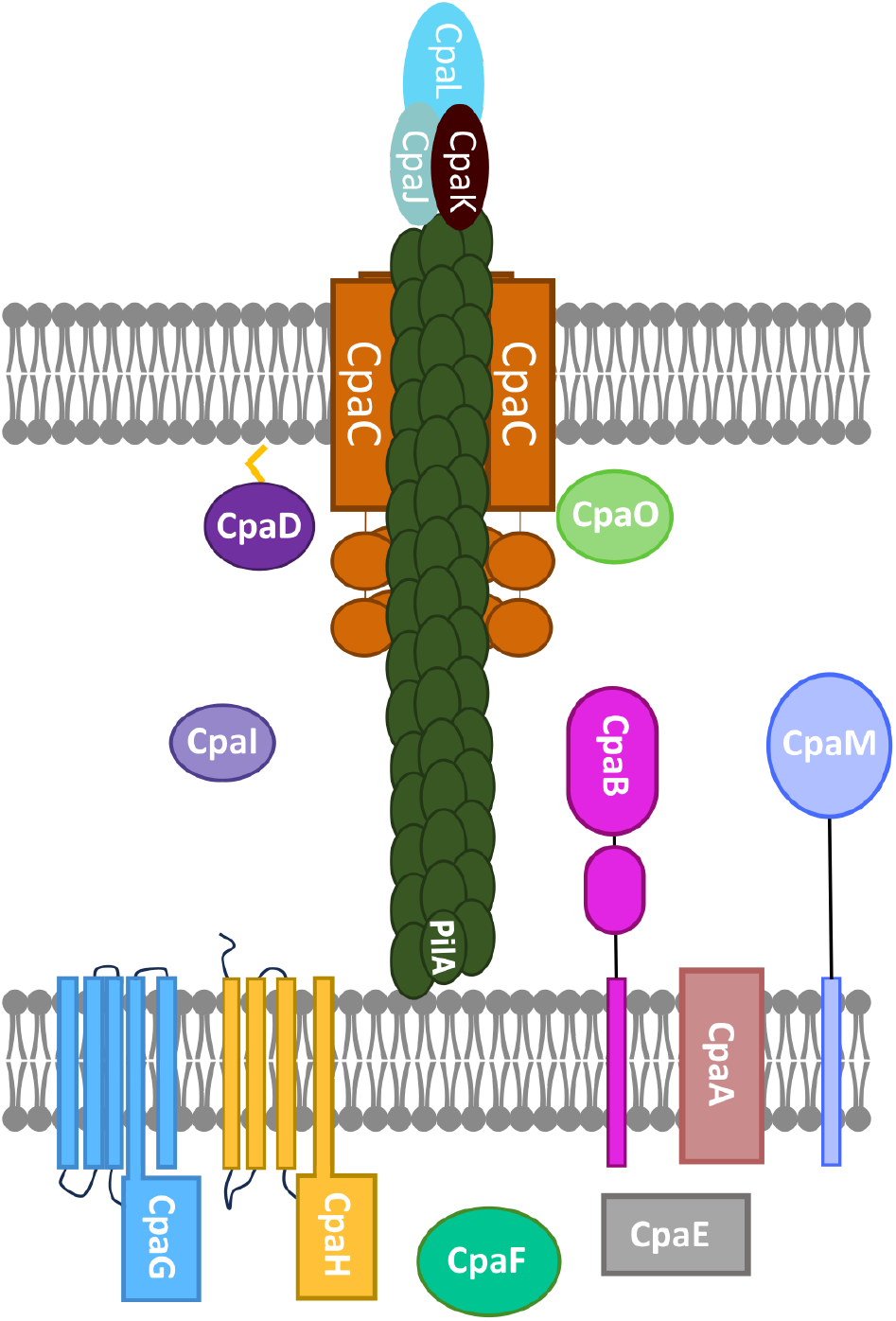
Schematic model of the *Caulobacter vibrioides* Tad pilus machine. For each Tad protein we predicted its domain organization, location, and function based on sequence and similarity to other type IV machines. The Tad pilus fiber is composed of the major pilin PilA (structure of the pilus PDB: 8UHF), and the tip is likely formed by CpaL, CpaJ, and CpaK. The Tad machine is constituted by the following proteins: CpaC forms the OM gate, CpaD is a lipoprotein that is likely the pilotin/secretin-associated protein, CpaO might be CpaC’s chaperone, CpaI is a periplasmic protein homologous to the CpaC N-terminus, CpaB is an inner-membrane-bound protein and is likely the alignment complex, CpaM is annotated as a polysaccharide deacetylase and could be involved in peptidoglycan remodeling to accommodate the Tad machine, CpaA is the prepilin peptidase, and CpaG and CpaH form the inner membrane platform. CpaF is the retraction and extension ATPase, and CpaE is involved in proper localization of the Tad machine to the cell pole.

### cpa *gene knockouts and ΦCbK resistance*

To investigate the role of *cpa* genes in Tad pilus production, we knocked out individual genes and tested each strain for ΦCbK resistance (Fig. S2). Our findings indicate that deletion of *cpaO* doesn’t confer resistance to ΦCbK infection and confirmed previous implications of *cpaI* importance for Tad assembly (Christen et al. 2016) using in-frame deletions and complementation assays.

### Fluorescence studies and Cpa assembly order

Previous fluorescence studies have shown that CpaC, CpaE, CpaF, and PleC each form polar foci, indicating precise co-localization and multimer formation of these Tad pilus machine components (Viollier et al. 2002; Ellison et al. 2019). These foci were associated with the leading pole for CpaC, CpaE, CpaF, and PleC, while CpaC also localized to the tip of the stalk. These studies also showed that the cytoplasmic protein CpaE is required for the modification and localization of the secretin CpaC; therefore, an inside-to-outside assembly model has been proposed.

To further investigate the Tad pilus machine assembly mechanism, we tested whether other Cpa proteins could form foci. Based on their predicted localization, we used mCherry C-terminally to CpaB (anchored in the inner membrane), CpaD (localized in the outer membrane), and CpaI (a predicted periplasmic protein). We inserted the sequences coding for these mCherry-fused variants into vanillate-inducible pMT335 plasmids and transformed the Δ*cpaB*, Δ*cpaD*, and Δ*cpaI* strains, respectively, with plasmids expressing each corresponding protein. We then imaged each strain for foci formation by epifluorescence microscopy.

The CpaB-mCherry and CpaD-mCherry strains formed bright fluorescent foci (Fig. 2A), but we didn’t detect any fluorescence in the CpaI-mCherry construct (data not shown). CpaB and CpaD foci localized at the leading pole, and CpaD formed additional foci at the tip of the stalk where CpaC was previously immunolocalized (Viollier et al. 2002). We next explored the ability of the CpaB-mCherry and CpaD-mCherry constructs to form foci in the presence of their wild-type alleles. We detected foci for both chimeric proteins with the same localization as the Δ*cpaB* and Δ*cpaD* strains (Fig. 2).

**Fig 2.**
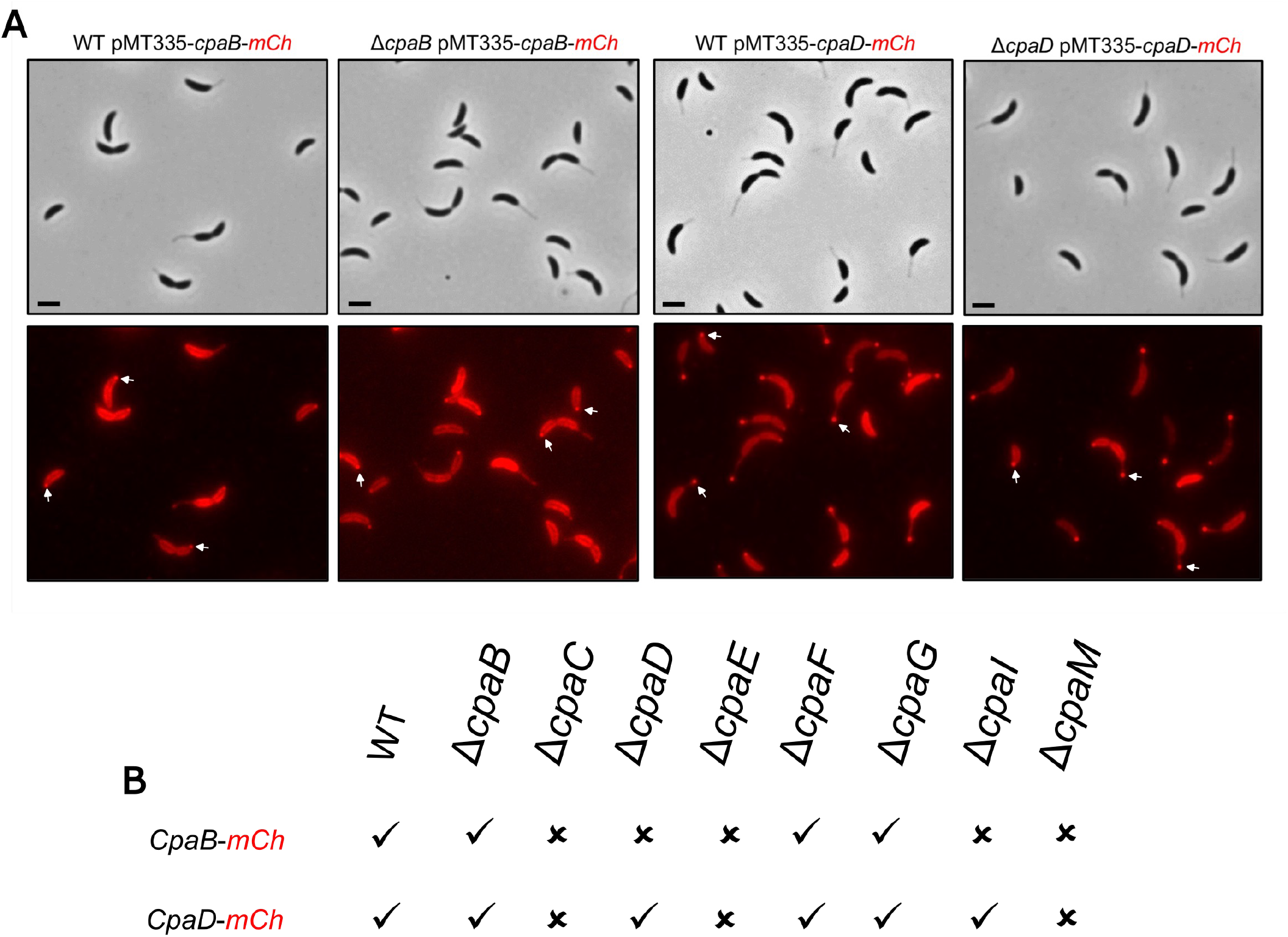
CpaB and CpaD foci formation and dependency. A) CpaB-mCherry and CpaD-mCherry strains form foci in absence or presence of the WT allele. Representative brightfield (top) and fluorescence (bottom) micrographs are shown (scale bar 5µm). B) Formation of CpaB-mCherry foci is dependent on the presence of CpaC, CpaD, CpaE, and CpaI, whereas CpaD-mCherry foci require CpaC, CpaE, and CpaM. These and other results suggest an assembly/dependency order of CpaM-CpaE, CpaC, CpaD, CpaI, then CpaB. (Representative brightfield and fluorescence micrographs are shown in Fig. S3-S4)

Based on this evidence, we investigated CpaB-mCherry and CpaD-mCherry foci formation in the WT, Δ*cpaB*, ΔcpaC, Δ*cpaD*, Δ*cpaE*, Δ*cpaF*, Δ*cpaG*, Δ*cpaI*, and Δ*cpaM* strains to dissect relative dependencies (results summarized in Fig. 2B, representative images Fig. S4, S5). Only the Δ*cpaC*, Δ*cpaE*, and Δ*cpaM* strains didn’t show CpaD foci, confirming CpaD dependency on CpaC, CpaE, and CpaM (the latter two being required for proper CpaC localization (Mignolet et al. 2016)) and showing CpaD assembly does not depend on CpaB, CpaF, CpaG, or CpaI. CpaB foci formation depends on CpaC, CpaD, CpaE, CpaI, and CpaM, but is independent of CpaF and CpaG.

These results provide new evidence that CpaI is involved in Tad pilus machine assembly and reveal that, like CpaC, CpaD also forms foci at the tip of the stalk in live cells. These results and others suggest an assembly order of CpaE > CpaM > CpaC > CpaD > CpaI > CpaB, which, except for beginning with the cytoplasmic protein CpaE, matches the outside-to-inside assembly pathway typical of other type IV machines (Friedrich et al. 2014).

### Tad architecture

To characterize the molecular architecture of the Tad pilus machine *in situ* we employed cryo-ET of various *C. vibrioides* mutants. We easily identified both piliated and empty machines in our tomograms (Fig. 3A–C). To resolve details, we aligned and averaged hundreds of Tad machines, aligning the periplasmic and cytoplasmic regions separately, as has been done previously (Chang et al. 2017).

**Fig 3.**
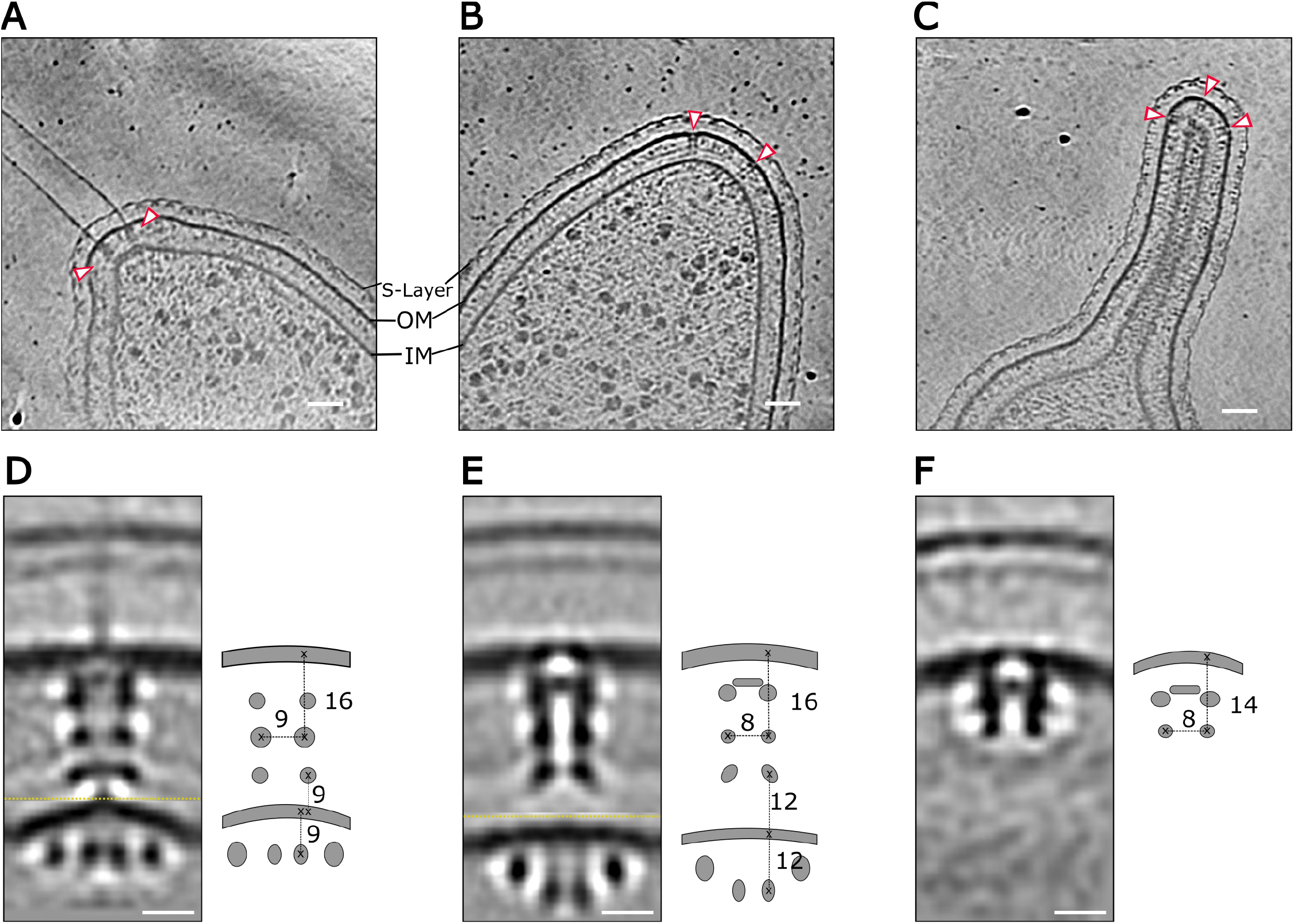
The Tad machine secretin and cytoplasmic rings exhibit three unique conformations. Representative tomographic slices showing (A) piliated machines, (B) non-piliated machines, and (C) secretins localized in the stalk (scale bar 200 nm). Composite subtomogram averages of the (D) piliated, (E) non-piliated, and (F) stalked states, with periplasmic and cytoplasmic halves (separated by yellow dashed lines) aligned separately with schematic representations showing distances (scale bars 10 nm).

The subtomogram average (STA) of the piliated machine exhibited three periplasmic rings (Fig. 3D) (OM–spanning pore, mid-periplasmic, and lower periplasmic) and two concentric cytoplasmic rings (inner cytoplasmic and outer cytoplasmic). The non-piliated STA (Fig. 3E) had the same overall architecture but with clear densities connecting the periplasmic rings to each other. The IM of the non-piliated machine was less bent than the piliated average, and its lower periplasmic ring was slightly farther from the IM (piliated, 9 nm, versus nonpiliated, 12 nm). Finally, the inner cytoplasmic ring of the non-piliated machine was substantially (~3 nm) farther from the IM, so the two cytoplasmic rings were no longer concentric.

We also found secretin-like densities at the tip of the stalk (Fig. 3C). The stalked STA was characterized by the presence of only an OM–spanning pore density and a small mid-periplasmic ring (Fig. 3F). In other type IV systems, the mid-periplasmic ring is formed by the interaction of the secretin’s N-terminal domain and the alignment complex (Chang et al. 2016; 2017; Korotkov et al. 2011), and a similar mechanism is likely present in the Tad system. Thus, our interpretation is that the OM–spanning pore density is formed by the CpaC-CpaD complex, confirming previous immunolocalization of CpaC to the tip of the stalk (Viollier et al. 2002).

### Localization of Tad components

To verify the position of each Tad protein in the STA envelopes, we collected tomograms of individual *cpa, pilA, podJ*, and *pleC* knockout strains. Because the absence of many of these genes prevents pilus synthesis, we focused our analysis on the non-piliated state, using the STA of the Δ*pilA* strain as our reference. In the Δ*cpaC*, Δ*cpaD*, Δ*cpaE*, Δ*cpaM, and* Δ*podJ* strains, we were unable to detect any Tad-machine-like densities. The STAs of the non-piliated states when Tad-machine-like densities are present are shown in Fig. 4 in order of completeness.

**Fig 4.**
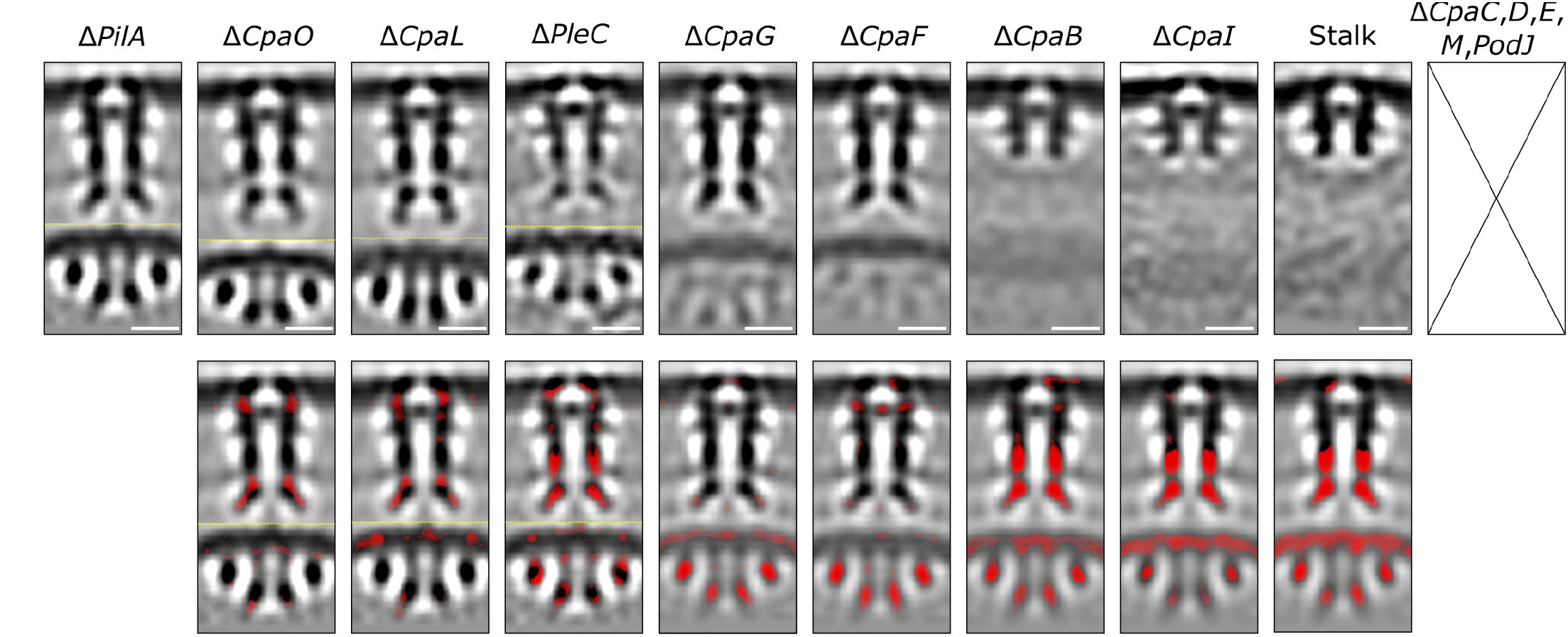
Subtomogram averages of deletion mutants. Top: Central slice of each knocked-out STA. Bottom: Difference maps (pilA-mutant) superimposed on the Δ*pilA* STA (red = density missing in the knocked-out strain) shown in order from most to least complete. No Tad machinery–like densities were found in the Δ*cpaC*, Δ*cpaD*, Δ*cpaE*, Δ*cpaM*, and Δ*podJ* strains (scale bars 10 nm).

The Δ*cpaO* and Δ*cpaL* strains had both piliated and non-piliated machines, though the piliated state occurred at a much lower frequency than in the WT (Fig. S6). The Δ*cpaO* and the Δ*cpaL* non-piliated STAs looked like the Δ*pilA* STA (Fig. 4) without any densities clearly missing, indicating that CpaO and CpaL are not core structural components of the machine, and are in fact dispensable for Tad machine assembly. The presence of the piliated Tad machine in Δ*cpaL* was not expected due to its ΦCbK resistant phenotype. During infection, the ΦCbK bacteriophage moves from the flagellum to the Tad pilus (Guerrero-Ferreira et al. 2011), which brings the phage in proximity of the S-layer. The presence of the piliated machine coupled with its ΦCbK resistance in the Δ*cpaL* strain indicates that CpaL is likely the ΦCbK phage receptor on the Tad pilus.

Given the importance of PleC in PilA accumulation and proper CpaE localization (Viollier 2002; Del Medico et al. 2020), we investigated Tad architecture in the Δ*pleC* mutant. We found only non-piliated machines at a low frequency (43 particles in 122 total tomograms compared to 181 particles in 183 tomograms for the Δ*pilA)*. The Δ*pleC* STA had a strong reduction of the mid-periplasmic, lower periplasmic, and outer cytoplasmic rings densities and a small reduction of the cytoplasmic ring, but all these were present, so PleC is not a core structural component of the machine. Given PleC’s location in the inner membrane with a bulky histidine kinase domain in the cytoplasm (Zhang et al. 2022) (Fig S1) and the results from the other mutants, we reasoned that the reduction of the outer cytoplasmic ring is a direct effect of PleC’s absence while the other ring reductions could be indirect effects.

In the Δ*cpaG*, Δ*cpaF*, Δ*cpaB*, and Δ*cpaI* strains, we found only non-piliated machines. The STAs of the Δ*cpaG and* Δ*cpaF* strains had no cytoplasmic densities, but their periplasmic portions were unaffected (Fig. 4). The STAs of the Δ*cpaB* and Δ*cpaI* strains closely resembled the stalked average, lacking the lower periplasmic ring, all cytoplasmic components, and part of the mid-periplasmic ring.

To precisely locate CpaB, CpaD, CpaE, CpaF, and PodJ inside our average map, we collected tomograms of fluorescently tagged proteins (mCherry or YFP) and subtracted each non-piliated STA from the Δ*pilA* reference STA (Fig. S7) to highlight any additional densities (Chang et al. 2016). Unfortunately, we could not detect any specific additional densities in the CpaD, CpaE, CpaF, or PodJ fluorescent strains. We speculate that in these cases, the mCherry was not rigidly attached, and therefore not visible in the averages. In the CpaB-mCherry strain, we detected a weak additional density in the lower periplasmic ring and the disappearance of the density connecting it to the mid-periplasmic ring. This perturbation could be the result of mCherry’s steric hindrance interfering with proper assembly, suggesting that CpaB forms the lower periplasmic ring.

Guided by the bioinformatic analyses, known structures, ΦCbK resistance phenotypes, fluorescence data, and cryo-ET maps, we concluded that CpaC and CpaD form the OM-spanning pore, CpaI likely forms part of the mid-periplasmic ring, CpaB forms the lower periplasmic ring, CpaG and CpaH form the IM complex, and CpaF forms the inner cytoplasmic ring (Fig. 5A). Because the outer cytoplasmic density showed a marked decrease in the Δ*pleC* strain, we speculate that the PleC-PodJ complex (Zhang et al. 2022) with the possible participation of CpaE forms the outer cytoplasmic ring, but we are unable to prove this due to the absence of Tad-machine-like densities in the Δ*podJ* and the Δ*cpaE* strains and lack of clear additional densities in the PodJ, CpaE, and CpaF YFP-tagged constructs.

**Fig 5.**
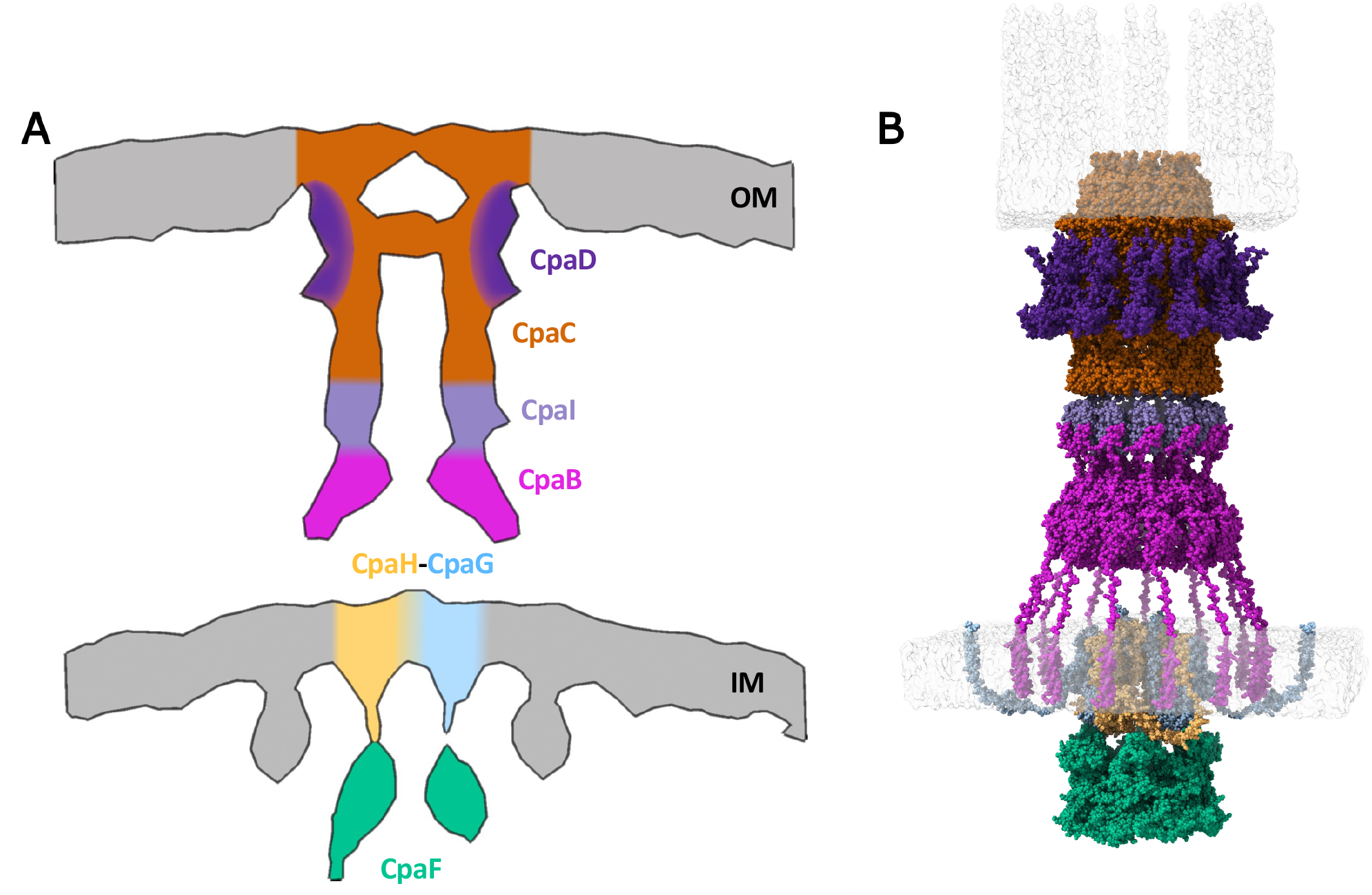
Tad architecture. (A) Localization of Cpa proteins in the Δ*pilA* STA based on fluorescence results and STAs of Δ*cpa* mutants. (B) Atomic representation of the integrative Tad machine model embedded in the bacteria membranes.

### Modeling

To gain more information about the Tad pilus machine assembly and architecture, we used ColabFold (Mirdita et al. 2022) to build an integrative model of the non-piliated Tad machine (Fig. 5B). For CpaC assembly, we used its AlphaFold predicted model in combination with the solved structure of a *Pseudomonas aeruginosa* homolog (Tassinari et al. 2023) (8ODN, Coverage: 31%, E-Value: 7e-34, Identity: 40.57%) to build a CpaC oligomer with C13 symmetry (the same symmetry as the *Pseudomonas* solved structure). Fitting the CpaC model into the stalked density revealed the presence of an empty flanking region (Fig. S8A) in our STA map. Based on our fluorescence data, which showed CpaD foci in the stalked cells and CpaD dependency on CpaC, we speculated that CpaD might occupy this empty region. We therefore generated a CpaC-CpaD model (Fig. S8) by using the CpaC C13 multimer as a scaffold to build a CpaC-CpaD multimer with C13 symmetry (Fig. S9). This model filled the empty space in the density map (Fig. S8B).

CpaB is composed of two domains that we believe form the lower periplasmic ring: the SAF and the RcpC (Fig. S1). The SAF domain is highly flexible (Iyer and Aravind 2004; Matsunami et al. 2016) and could be responsible for the high structural flexibility revealed in the Δ*cpaO* and Δ*cpaL* STAs (Fig. 4). Assuming a one-to-one molar ratio between the secretin and the lower periplasmic ring (Chang et al. 2016), we built the CpaB ring with the same C13 symmetry as the CpaC-CpaD complex. We started by building the CpaB_141-297_ C13 assembly, then we rigid-body fit it to the lower periplasmic ring (Fig. S10). We then built CpaB_48-141_ using the CpaB monomer AlphaFold model (Jumper et al. 2021), which we also used to insert the transmembrane helix CpaB_1-29_ into the IM density. We finished the CpaB model by building the flexible loop (CpaB_30-47_) with Modeller (Šali and Blundell 1993).

Our results suggested that CpaI is an adaptor protein that links CpaC to CpaB, so we built a model of the CpaB-CpaI-CpaC interface (Fig. S11). In the model, the two C-terminal β-strands of CpaB bind CpaI and, in a similar way, the last two CpaI β-strands contact CpaC. It should be noted that in the usher pilus, the pilin subunits use N-terminal extensions, which are not part of their Ig-like fold, for polymerization (Geibel and Waksman 2014); therefore, a similar mechanism could be used by the Tad system for protein-protein interaction. Given the low resolution of the STA map and the low ipTM score of the CpaB-CpaI-CpaC multimer, our confidence in this region is not as high as the previously described portions. However, a recent study by Evans and collaborators (Evans et al. 2024) confirmed that in *Pseudomonas aeruginosa*, the two C-terminal β-strands of CpaB are necessary for Tad pilus function, increasing confidence in our model. Moreover, our model can explain the lack of the CpaI density in the Δ*cpaB* strain: CpaI might be held in place by CpaC and CpaB via two flexible loops, and the loss of CpaB frees CpaI from one of the two restraints, allowing the protein to move more freely, which leads to its disappearance in averages.

The IM-platform proteins CpaG and CpaH were modeled based on their similarities with archaeal proteins. CpaG and CpaH derived from a gene fission event in Archaea, and the combined CpaG-CpaH topology matches that of the archaellum IM-platform protein FlaJ (Denise et al. 2019). Recently, the FlaJs of *Sulfolobus acidocaldarius* (*Sa*) and *Pyrococcus furiosus* were modeled as dimers, which is in line with the type IV IM-platform homolog PilC (Bischof et al. 2016; Nuno de Sousa Machado et al. 2022). We therefore built the CpaG-CpaH complex as a tetramer (Fig. S12A-D). In addition to the AlphaFold scores, we tested the quality of the model by inserting it into a membrane using PPM 3.0 (Lomize et al. 2022). In this model all the transmembrane helices were in the lipid bilayer (Fig. S12E), which, combined with the analogies to Archaea, increases our confidence in the model.

We built the model of the bifunctional ATPase CpaF as a hexamer according to its solved structure (Hohl et al. 2024) (Fig. S13). Because the localization of CpaF depends on CpaG’s presence, we reasoned that the CpaF-CpaG-CpaH interaction is similar to that of *Sa*FlaJ-FlaI, which has been shown to depend on the positively charged cytoplasmic loops of *Sa*FlaJ and the negatively charged N-terminal crown domain of *Sa*FlaI (Reindl et al. 2013). We therefore investigated these regions in Alphaproteobacteria and found that the charge of the linker connecting the first and second transmembrane helices of CpaG and CpaH has a conserved positive charge at neutral pH, with isoelectric points between 10–12 and 9.1–11.5, respectively (Tables S1, S2). In addition, the electrostatic potential of the CpaF crown domain resembles that of FlaI (Fig. S14). It is therefore likely that CpaF interacts with CpaG-CpaH through a *Sa*FlaJ-FlaI-like mechanism.

### Tad assembly mechanism

Many type IV machines and type II secretion systems have been reported to assemble in an outside-to-inside pathway (Friedrich et al. 2014; Korotkov et al. 2012). In *Caulobacter*, however, the formation of cytoplasmic localization scaffolds (PodJ-CpaE-PleC and ZitP-CpaM) is the first step in Tad machine assembly and, therefore, a mixed inside-to-outside assembly mechanism has been hypothesized (Tomich et al. 2007; Mignolet et al. 2018).

Our fluorescence and cryo-ET data support such a mixed inside-to-outside assembly mechanism. By imaging Tad machines in mutants, we captured snapshots of the assembly process at different stages (Fig. 6). Our cryo-ET data showed that the first part of the machine to assemble is the OM-spanning pore formed by the CpaC-CpaD complex. This complex depends on the cytoplasmic localization scaffolds, which means a signal must somehow be transmitted from the cytoplasm to the outer membrane. Our bioinformatics analysis points to PodJ as the probable transmitter, and the fluorescence and cryo-ET data reveal CpaE as a necessary accessory protein. The absence of CpaD leads to a lack of the OM-spanning pore density, which indicates that CpaD helps secretin assembly, which also requires CpaM. After the formation of the CpaC-CpaD complex, CpaI likely binds to CpaC’s N-terminal region and helps recruit the CpaB ring, thus completing the assembly of the periplasmic portion of the machine.

**Figure 6.**
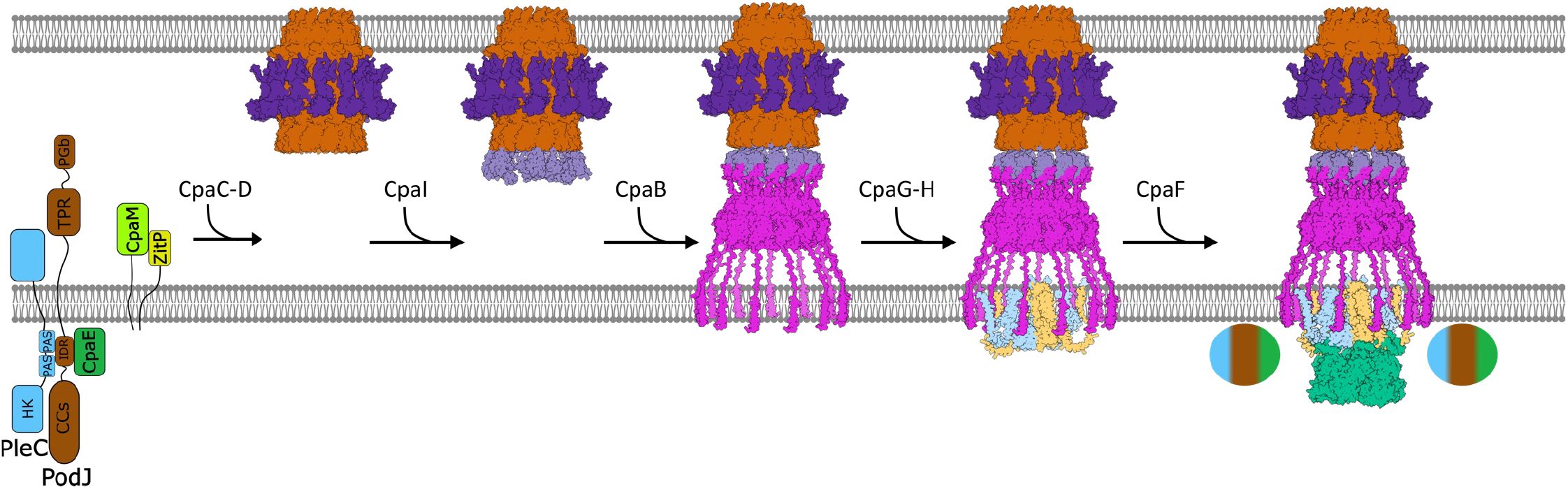
Tad machine assembly model. Our results suggest a stepwise assembly mechanism for the Tad machinery. Assembly begins with the formation of PleC-PodJ–CpaE and ZitP–CpaM platforms at the new cell pole, promoting CpaC–CpaD complex formation. CpaI then binds CpaC and recruits CpaB, completing the periplasmic components. Subsequently, the CpaG–CpaH complex forms in the inner membrane, recruits CpaF, and stabilizes the unresolved cytoplasmic outer ring. Interestingly, in stalked cells, the secretin and OM structures observed disappear when CpaF is deleted, but they remain when other last recruits (CpaG, CpaH) are removed. Therefore, we cannot exclude the possibility that CpaF may perform a function necessary for the formation of the secretin complex at the pole and tip of the stalk.

Deletion of CpaG and CpaF led to the disappearance of both cytoplasmic rings, suggesting that a complex protein-protein network could be formed by the CpaG-CpaH-CpaF complex and the proteins that form the outer cytoplasmic ring. In Archaea, CpaF homologs interact with FlaH, an ATP-binding protein with an HMM profile resembling CpaE (Denise et al. 2019). Therefore, CpaE alone or in combination with PodJ and PleC might form the outer cytoplasmic ring. These three proteins could be present at the base of the Tad machine in early assembly stages, stabilized as a ring only in the presence of all cytosolic components.

Interestingly, in stalked cells, the secretin and OM structures observed disappear when CpaF is deleted, but they remain when other last recruits (CpaG, CpaH) are removed. Therefore, we cannot exclude the possibility that CpaF may perform a function necessary for the formation of the secretin complex at the pole and tip of the stalk.

## DISCUSSION

The *Caulobacter vibrioides* Tad pilus system is a peculiar type IV machine because of its unique spatiotemporal regulation and its position in the type IV filament family evolutionary tree. Here, we combined fluorescence microscopy, cryo-ET, subtomogram averaging (STA), and modeling to characterize the architecture and order of assembly of the Tad machine.

Several proteins involved in Tad production (CpaC, CpaE, CpaF, and PleC) form bright foci when tagged with fluorescent proteins (Viollier et al. 2002; Ellison et al. 2019). In the present work we have expanded this list to include CpaB and CpaD. Since CpaB and CpaD are predicted to be in the two boundaries of the Tad periplasmic portion (CpaB is predicted to be an inner membrane protein and CpaD an outer membrane lipoprotein) and they both form foci in the presence of the WT allele, we were able to investigate the assembly order using several *cpa* knocked-out strains. Based on our results the secretin-periplasmic subcomplex assembles with a CpaC ≥ CpaD > CpaI > CpaB, with CpaB showing a strong dependency on CpaI.

The *Caulobacter vibrioides* Tad pilus is the first type IV filament family member that shows the ATPase density in the non-piliated state. Since CpaF is responsible for both extension and retraction, the increased distance between the inner cytoplasmic ring and the IM in the non-piliated state (9 vs 12 nm) might be the result of a clutch-like mechanism disengaging the ATPase from the IM platform when not actively involved in extension/retraction.

Investigation of the protein identities of the cytoplasmic densities was complicated by the complete absence of machines in both PodJ and CpaE deletion mutants. Moreover, deletion of CpaF or CpaG led to the absence of both cytoplasmic rings. The best candidate to occupy this region is the PleC-PodJ-CpaE scaffold given its role in the polar positioning of the Tad pilus (CpaE is a distant homolog of archaeal proteins arCOG00589 and arCOG05608 involved in archaellum formation).

The absence of PleC slightly reduced both cytoplasmic densities and destabilized the CpaB and CpaI rings, leading to the speculation that the outer cytoplasmic ring might be formed by all three proteins. Interestingly, the periplasmic portion of PodJ has been shown to be involved in pili formation (Viollier et al. 2002) and consists of a PG binding domain and a TPR domain, the latter of which appears to be involved in secretin stabilization (Silva et al. 2020). The *Pseudomonas aeruginosa* TPR-containing protein CpaO acts as a chaperone for CpaC assembly, and the two protein structures have been solved in complex by cryo-EM (Tassinari et al. 2023). In all our experimental conditions, the Δ*cpaO* strain had a functional Tad machine; therefore, it is possible that PodJ’s TPR domain is CpaC’s chaperone. Soon after a new flagellated cell is born, PodJ’s periplasmic domain is cleaved (Viollier et al. 2002), which would prevent assembly of new secretins, providing both temporal and spatial control over Tad biogenesis.

## METHODS

### Bacterial strains, mutant generation, growth conditions, and bacteriophage ΦCbK infection assays

The *C. vibrioides* NA1000 565050 wild-type cells were used in this study. All *C. vibrioides* strains used in this study were derived from NA1000. All knockout and tagged mutants were either from J. M. Skerker (2000) or generated herein.

Growth conditions were adapted from (Ely 1991) to enrich for swarmer cells that exhibit piliation at their cell poles. Single colonies were inoculated overnight from a 2% M2G agar plate into 5 mL of M2G media incubated at 30 °C at 220 rpm. Each 5 mL overnight culture was backdiluted in 15 mL of M2G and grown at 30 °C at 220 rpm for 3 hours to an OD_600_ of 0.7. The cells were then transferred to a conical centrifuge tube and centrifuged at 5,525 x g for 20 minutes. The supernatant was aspirated, and the remaining pellet was resuspended with 1 mL of cold M2 media and transferred to a 2-mL microcentrifuge tube and centrifuged at 15,000 x g for 3 minutes. The supernatant was aspirated, and the remaining pellet was resuspended with 900 µL of cold M2 media and mixed with 900 *µ*L of cold Percoll^®^ solution (Sigma-Aldrich #P4937). The mixture was then centrifuged at 15,000 x g for 20 minutes. After the fractionation, the supernatant along with the top band of stalked cells were discarded, and the remaining bottom band of swarmer cells was washed three times with 1 mL of cold M2. The cell pellet was resuspended in 20 *µ*L of M2 for imaging.

The plasmid pRVMCS-5_spmX-mCherry (Radhakrishnan et al. 2008) was digested with EcoRI and XbaI to isolate the coding sequence of mCherry. This fragment was then inserted into the multiple cloning site (MCS) of plasmid pMT335, previously digested with the same enzymes. Next, the genes *cpaB* and *cpaD* (without the stop codon) were amplified using primer pairs containing NdeI and EcoRI restriction sites and subsequently cloned into this vector. This resulted in C-terminal fusions of the proteins with mCherry. Below is the EcoRI–mCherry–XbaI DNA sequence cloned into pMT335:

>EcoRI–mCherry–XbaI gaattcgggatccacggtggtggtggtggtgtcgtgtcgaagggtgaagaagataatatggccatcatcaaggagtttatgcgcttcaaggtccacatggagggctcggtcaacgggcacgagttcgaaatcgagggcgagggcgaaggtcgcccgtatgagggcacccagaccgccaagctgaaggtgaccaagggcgggcccctgccgttcgcgtgggacatcctgtcgccgcagttcatgtatggttcgaaggcctatgtgaagcacccggcggacatcccggactacctgaagctctccttcccggagggctttaagtgggagcgggtgatgaacttcgaagatggtggcgtcgtcacggtcacccaggacagctcgctgcaggacggcgagttcatctacaaggtcaagctgcgcggcacgaacttcccgtccgatggcccggtcatgcagaagaagacgatgggctgggaggcgtcgtccgaacggatgtatcccgaggacggggccctgaagggcgagatcaagcagcgcctcaagctgaaggacggcggccactacgatgccgaggtgaagacgacctacaaggccaagaagcccgtccagctgcccggggcctacaacgtcaacatcaagctggacatcacctcccacaacgaagattatacgatcgtggagcagtatgagcgcgccgagggccgccactcgaccggcggcatggatgagctctacaagtaatctagctgcagcccgggggatccactagttctaga >mcherry EFGIHGGGGGVVSKGEEDNMAIIKEFMRFKVHMEGSVNGHEFEIEGEGE GRPYEGTQTAKLKVTKGGPLPFAWDILSPQFMYGSKAYVKHPADIPDYLK LSFPEGFKWERVMNFEDGGVVTVTQDSSLQDGEFIYKVKLRGTNFPSDGP VMQKKTMGWEASSERMYPEDGALKGEIKQRLKLKDGGHYDAEVKTTYKAK KPVQLPGAYNVNIKLDITSHNEDYTIVEQYERAEGRHSTGGMDELYK

### Cryo-ET sample preparation and fast-incremental single exposure (FISE) imaging

The Au solution used for cryo-ET fiducial tracking was prepared by mixing 1 mL of 10-nm gold colloidal beads (Sigma-Aldrich) with 250 *µ*L of 5% BSA in PBS. The mixture was then vortexed and centrifuged at 18,000 x g for 30 minutes. The final supernatant aspiration left 20 *µ*L of resuspended Au solution to be mixed with 20 *µ*L of resuspended cell solution in a 1:1 ratio. R3.5/1 or R2/2 carbon coated 200 mesh copper Quantifoil grids (Quantifoil Micro Tools, GmbH, Jena, Germany) were glow discharged for 60 seconds to prepare hydrophilic surfaces before sample application. The mixed cell solution was added to the EM grids at 3 *µ*L and plunge-frozen using a Vitrobot Mark IV (FEI Company, Hillsboro, OR) with 100% humidity. A blot force setting of 2 and blot time setting of 4 seconds were applied with the extra fluid blotted off using Whatman filter paper. The grids were then plunge-frozen in a liquid ethane/propane mixture.

All samples were imaged using a 300 keV Titan Krios (Thermo Fisher Scientific) electron cryo-microscope with a K3 Summit direct detector, energy filter (Gatan, Pleasonton, CA), and high-precision stage for fast tilt-series acquisition. Tilt series were collected using the FISE scheme, with a constant angular increment of 3° from −60 to 60 degrees. A dosage of 180 e^−^/Å^2^ was applied with a defocus of −8 *µ*m, pixel size of 3.3 Å, and magnification of 26,000X. SerialEM software (Mastronarde 2005) was used for sample acquisition and a RAPTOR software pipeline (Ding et al. 2015) was used to create three-dimensional reconstructions of tilt series (Chreifi et al. 2019; 2021; Eisenstein et al. 2019). Representative tomograms were denoised using ISONET (Liu et al. 2022).

### Subtomogram averaging and difference analysis

The basal bodies of the Tad machine were identified within the cell envelopes at the cell pole by visual inspection. Subtomogram averaging was performed using the IMOD PEET sub-volume averaging program with 2-fold symmetrization along the y-axis (Nicastro et al. 2006). All the final averages were realigned using the Δ*pilA* strain as a reference in PEET, and the central slice of each volume was saved as an image. Difference analysis was conducted by creating difference maps in GIMP (Wilber 2019).

## Acknowledgements

This work was funded by the National Institutes of Health grant R01 AI127401 to G.J.J. Cryo-ET data collection was performed at the Beckman Institute Resource Center for Transmission Electron Microscopy at the California Institute of Technology. We thank Dr. Jane Ding and Dr. Songye Chen for valuable discussions on optimal cryo-ET data collection and processing. We are grateful to Prof. Yi-Wei Chang (University of Pennsylvania), Dr. Stefan Kreida, Dr. Catherine M. Oikonomou, and Rachel Webb for their critical feedback and editorial assistance.

## Author contributions

S.M. and R.R. prepared the samples, performed the cryo-ET data collection, and analyzed the tomography data. S.M. performed bioinformatics analysis and pseudoatomic modeling. G.P. and P.V. generated the *C. vibrioides* mutants and ϕCbK infection assays. P.V. and G.J.J. conceptualized the project and analyzed the data. R.R., S.M., G.P., P.V. and G.J.J. wrote the manuscript.

