## Supplementary Figures for "Electron Cryo-Tomography Reveals the *Caulobacter vibrioides* Tight Adherence Pilus Architecture"

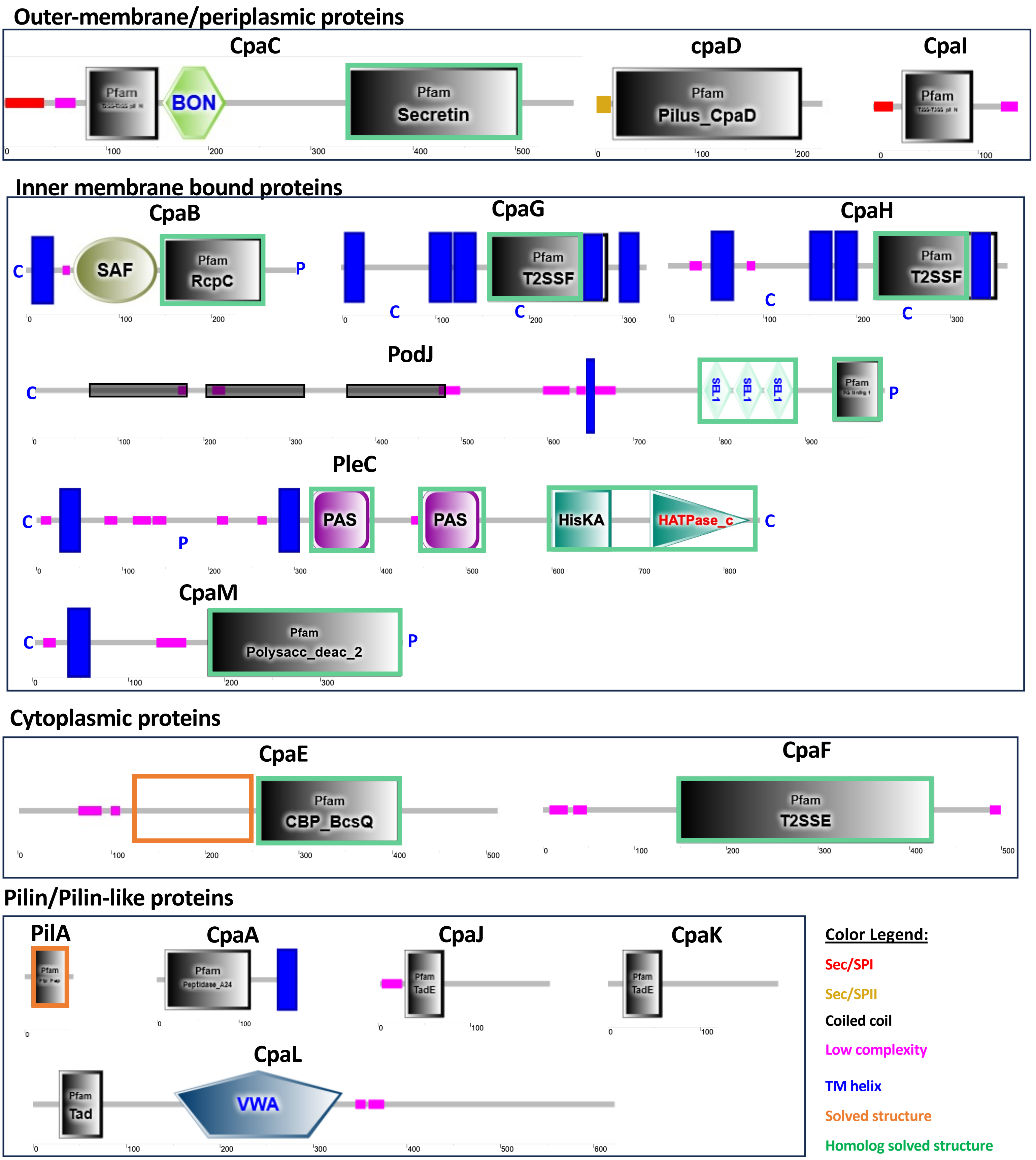


**Fig S1. | Domain architecture of Cpa proteins.** Proteins are grouped by their predicted localization and functions. CpaC, CpaD, and CpaI show a Sec/SPI or Sec/SPII signal peptide. CpaB, CpaG, CpaH, PodJ, PleC, and CpaM are predicted to be inner-membrane bound (C=Cytoplasm, P=Periplasm). CpaF and CpaE are cytoplasmic proteins. PilA is the major pilin, CpaA is the PilA prepilin peptidase, and CpaJ, CpaK, and CpaL are minor pili (Data collected from the SMART database).

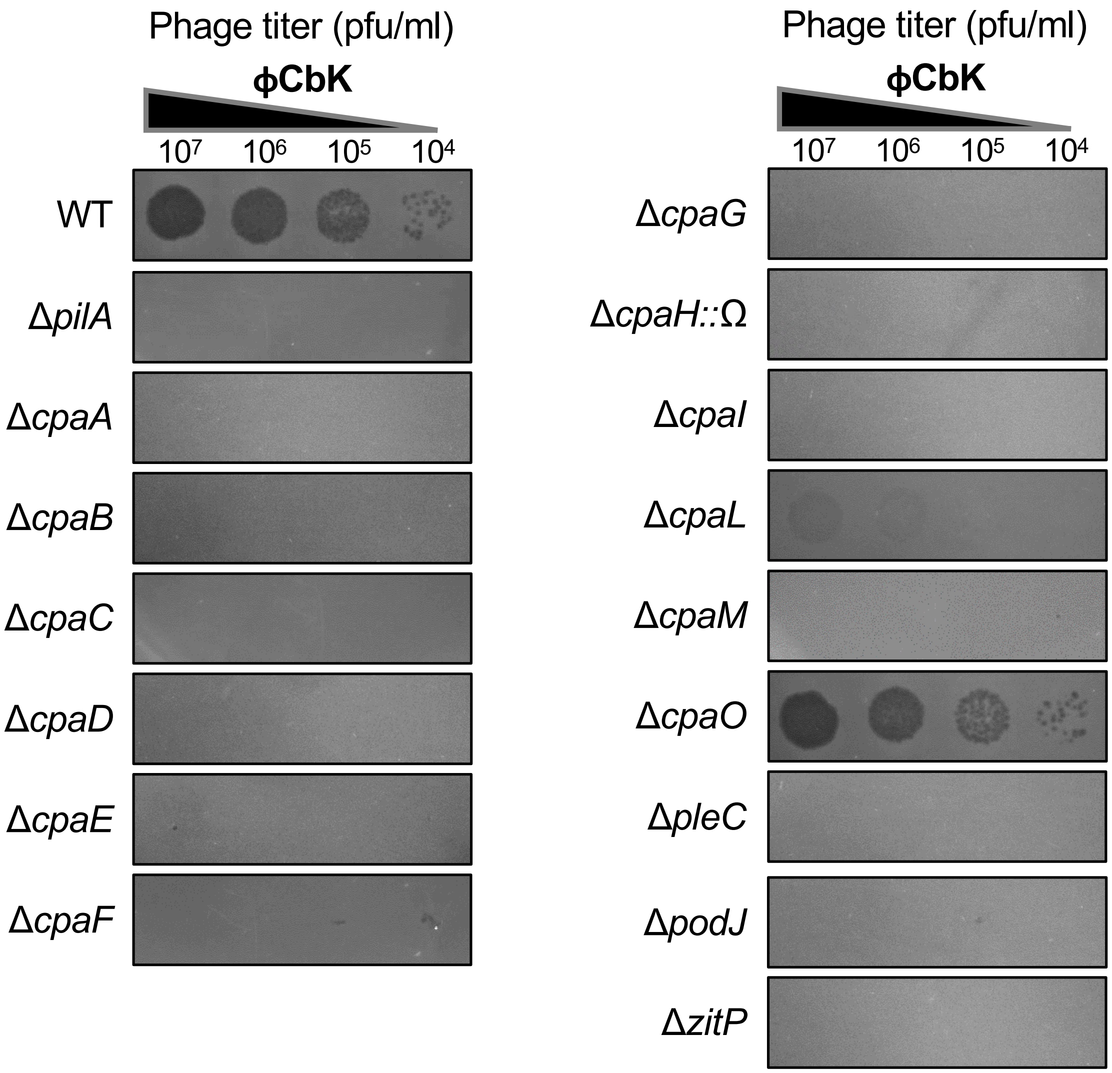


**Fig S2. | ϕCbK infection screening.** Knocked-out strains of *cpa* and other genes involved in Tad polar localization were screened for plaque formation after infection with ϕCbK. All are essential for ϕCbK infection (and by inference, Tad pilus biogenesis and function) except *cpaO*. *cpaK* and *cpaJ* were not tested.

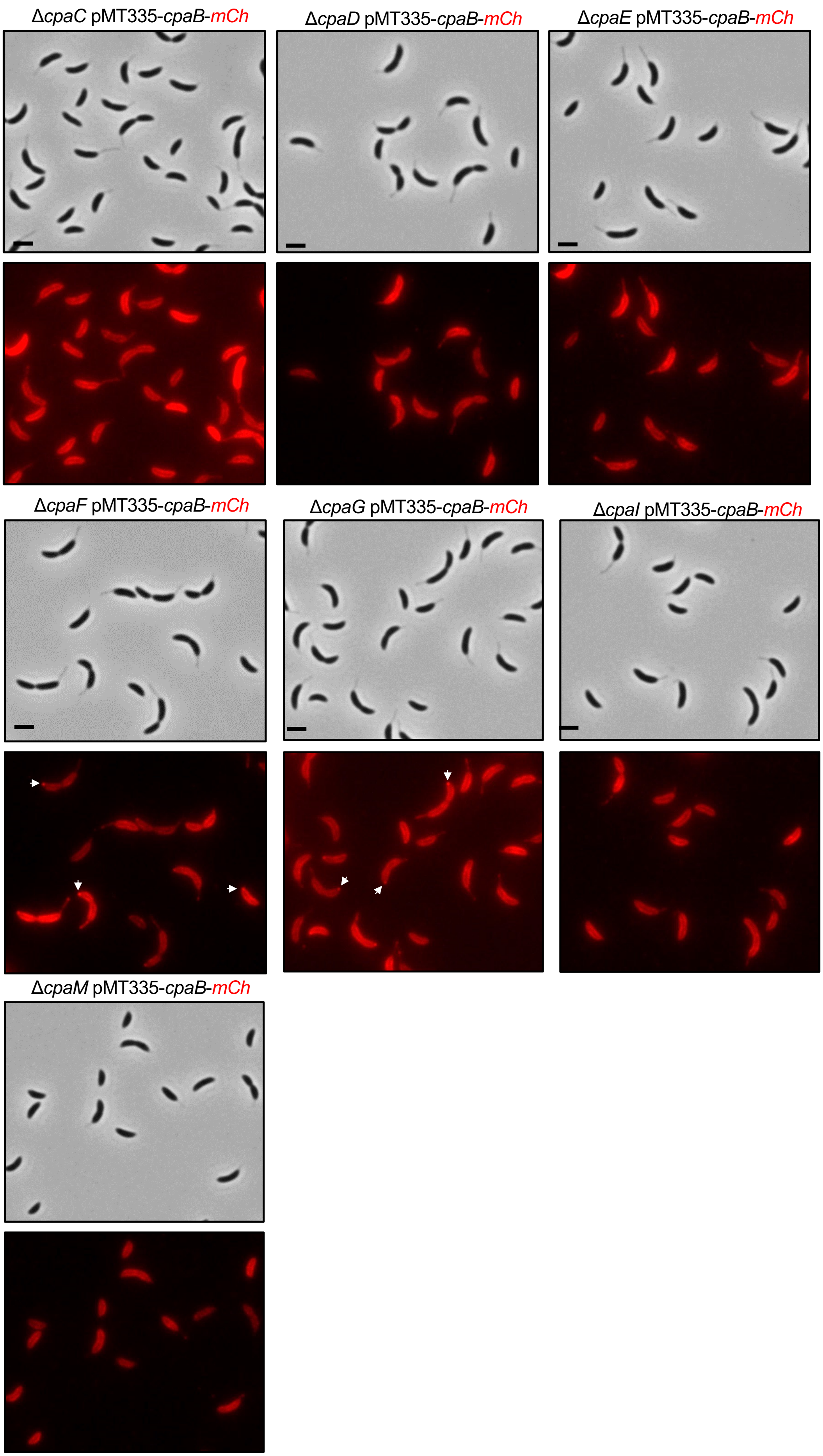


**Fig S3. | Representative brightfield and fluorescence micrographs of CpaB-mCherry strains.** CpaB-mCherry forms foci in the Δ*cpaF* and Δ*cpaG* strains.

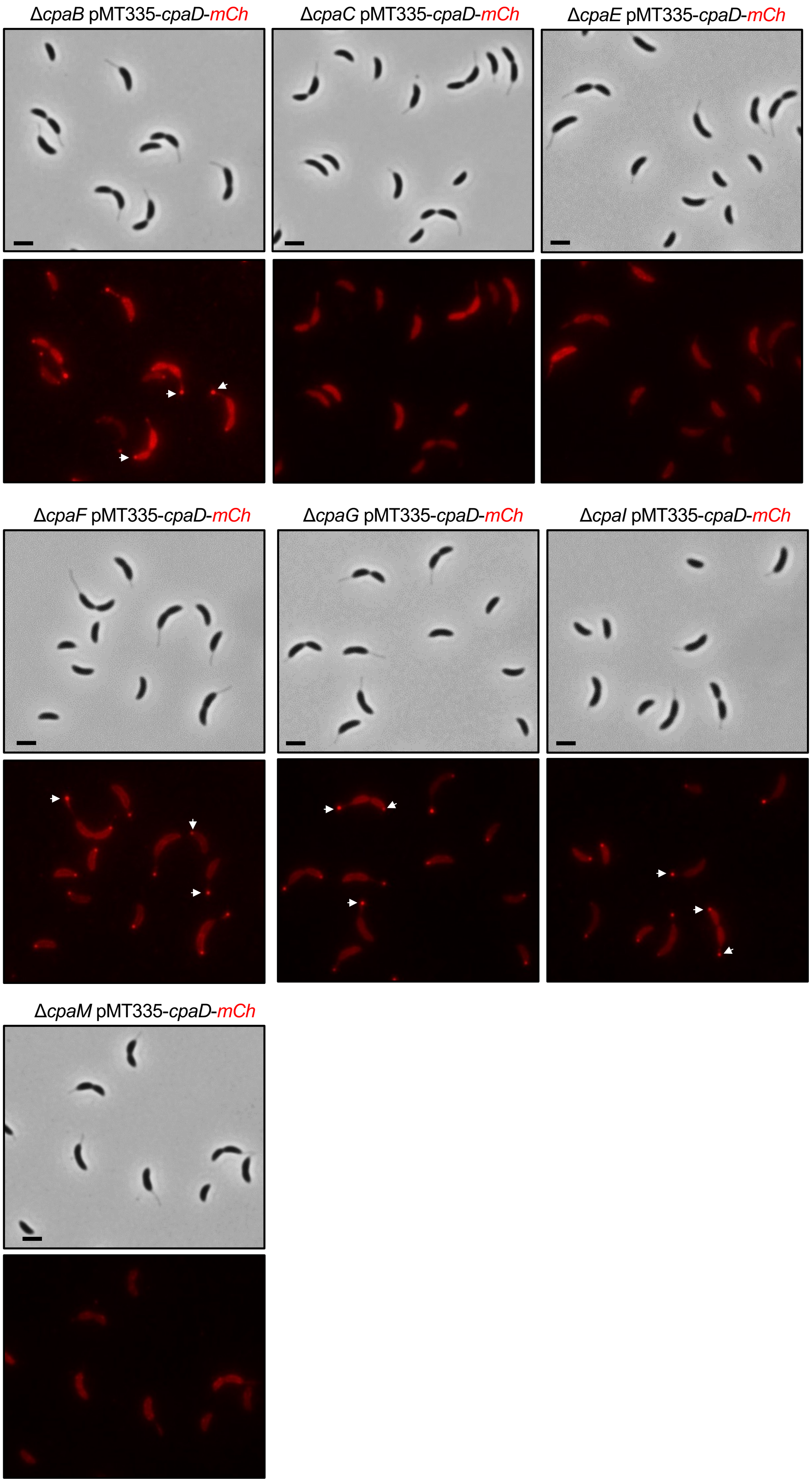


**Fig S4. | Representative brightfield and fluorescence micrographs of CpaD-mCherry**. CpaD-mCherry forms foci in Δ*cpaB*, Δ*cpaF*, Δ*cpaG*, *and* Δ*cpaI* strains.

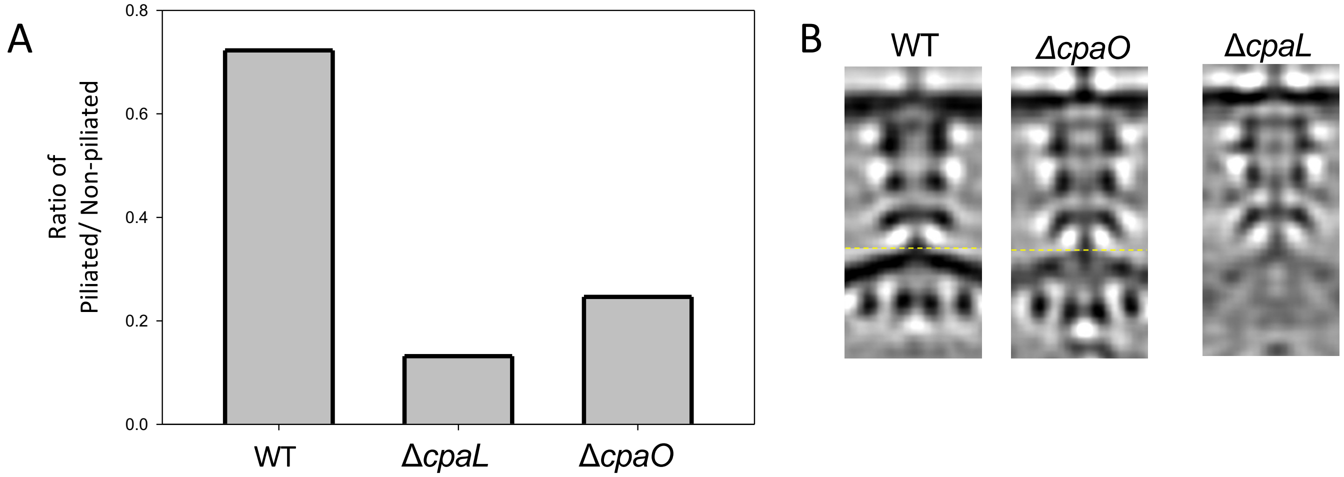


**Fig S5. | Δ*cpaL* and Δ*cpaO* STA**. A) Histogram of the ratio between the piliated and non-piliated Tad machines found in the WT, Δ*cpaL*, and Δ*cpaO* strains. B) STAs of piliated WT, Δ*cpaO*, and Δ*cpaL* strains.

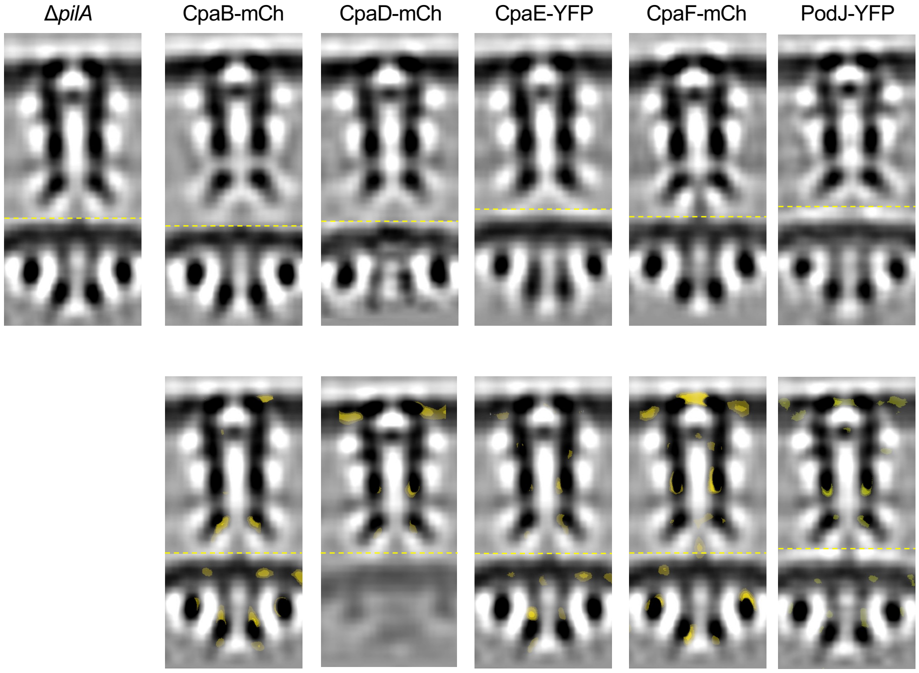


**Fig S6. | Subtomogram averages of *cpa* fusion mutants.** Top: Central slice of the subtomogram averages of the non-piliated machine in Δ*pilA,* CpaB-mCh, CpaD-mCh, CpaE-YFP, CpaF-mCh, and PodJ-YFP Δ*cpaM* strains (dotted line indicates the stitching position). Bottom: Differences superimposed on Δ*pilA* map (yellow = additional densities).

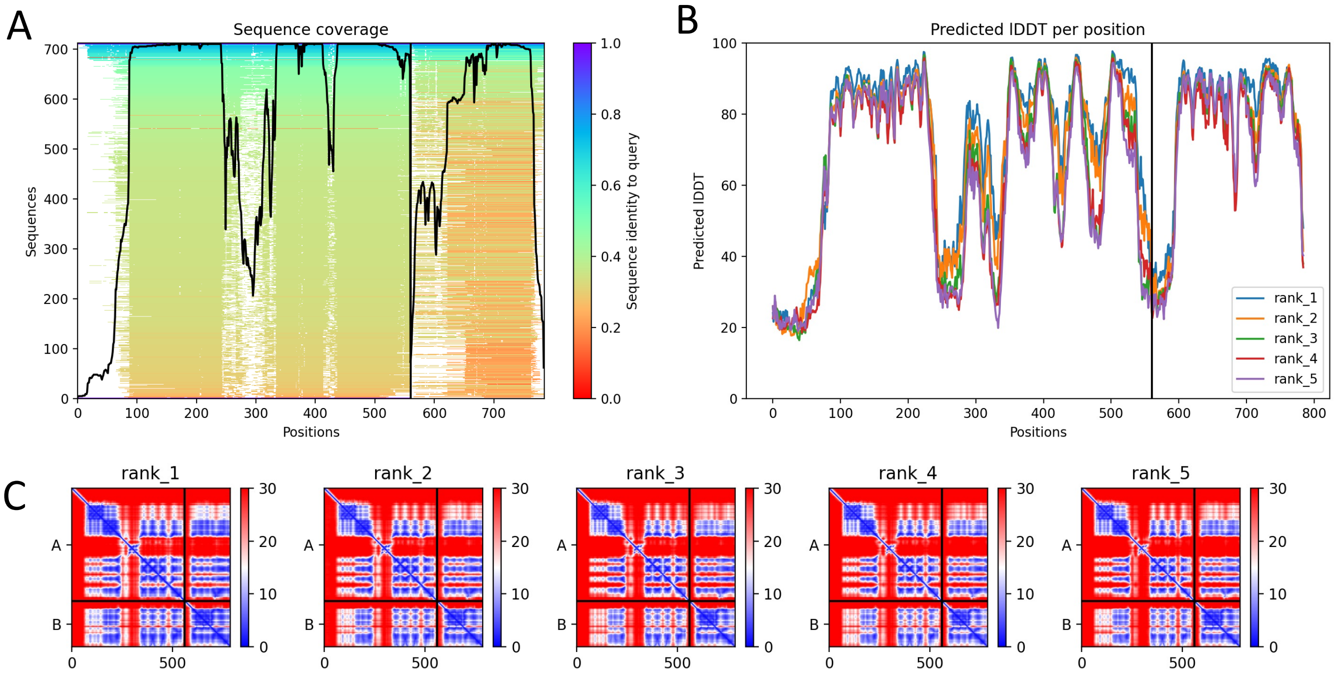


| | pLDDT | pTM | ipTM |
|---|---|---|---|
| model1 | 66.7 | 0.598 | 0.749 |
| model2 | 65.2 | 0.59 | 0.736 |
| model3 | 73.6 | 0.643 | 0.785 |
| model4 | 70.9 | 0.626 | 0.76 |
| model5 | 64.7 | 0.587 | 0.736 |

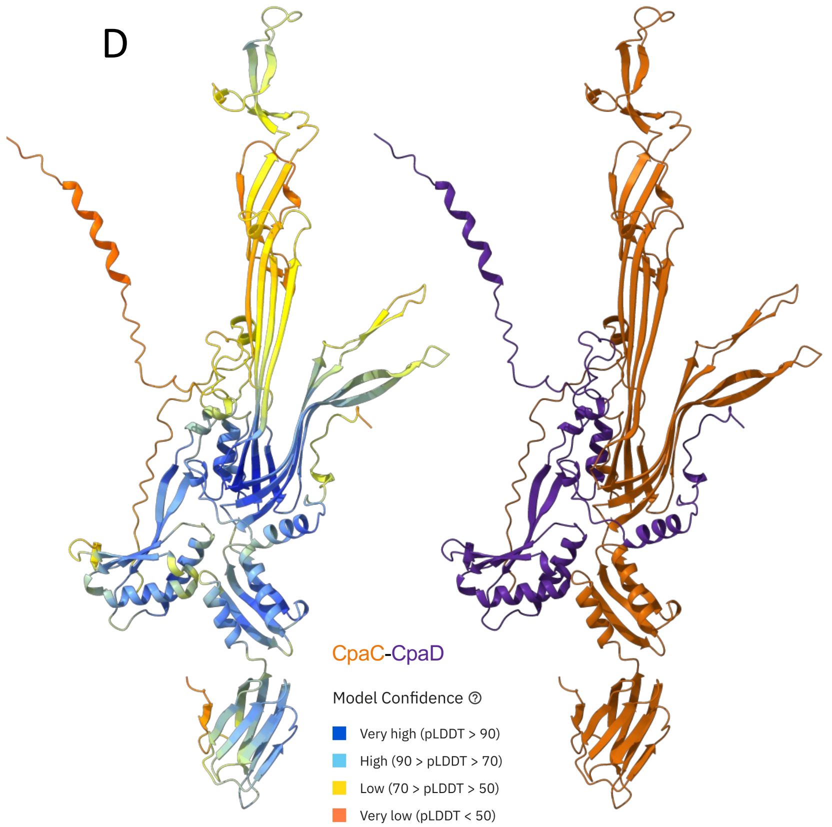


**Fig S7. | CpaC-CpaD AlphaFold model.** (A) Sequence coverage, (B) pLDDT, (C) PAE, and table with metrics of the five different models for the CpaC-CpaD heterodimer. (D) Best model colored by confidence and by chain identity.

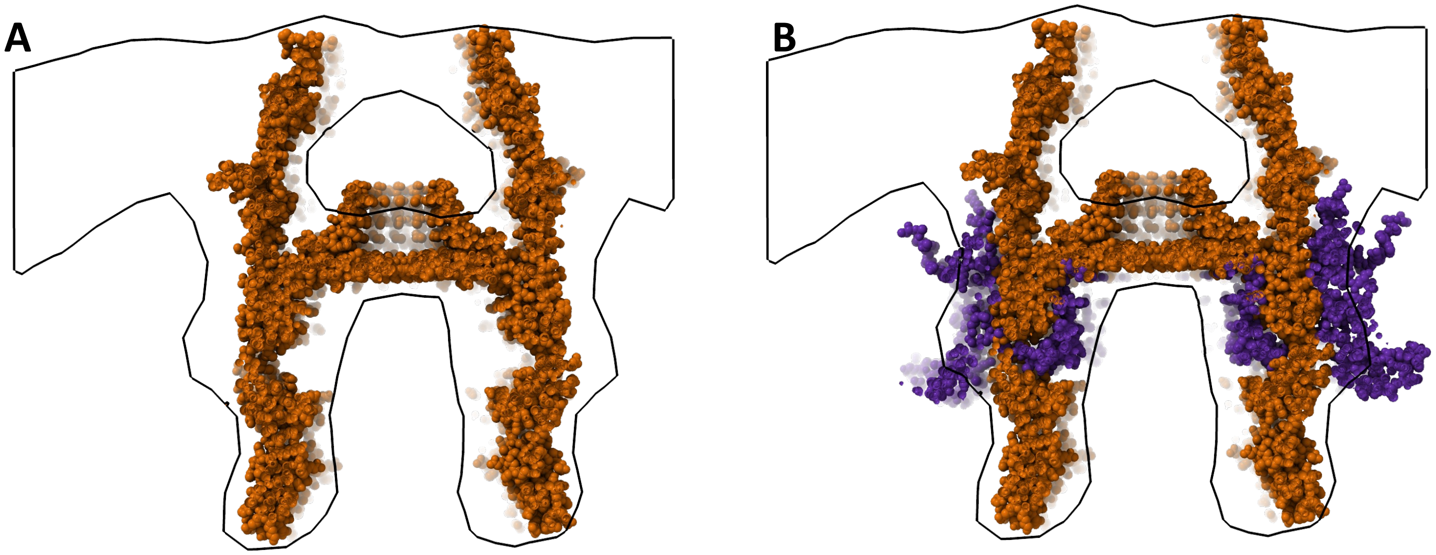


**Fig S8. | CpaC and CpaC-CpaD models fit into the stalked STA density.** Fitting of the (A) C13 CpaC model and (B) the C13 CpaC-CpaD model in the Tad stalked STA. The CpaC-CpaD model better fills the density envelope.

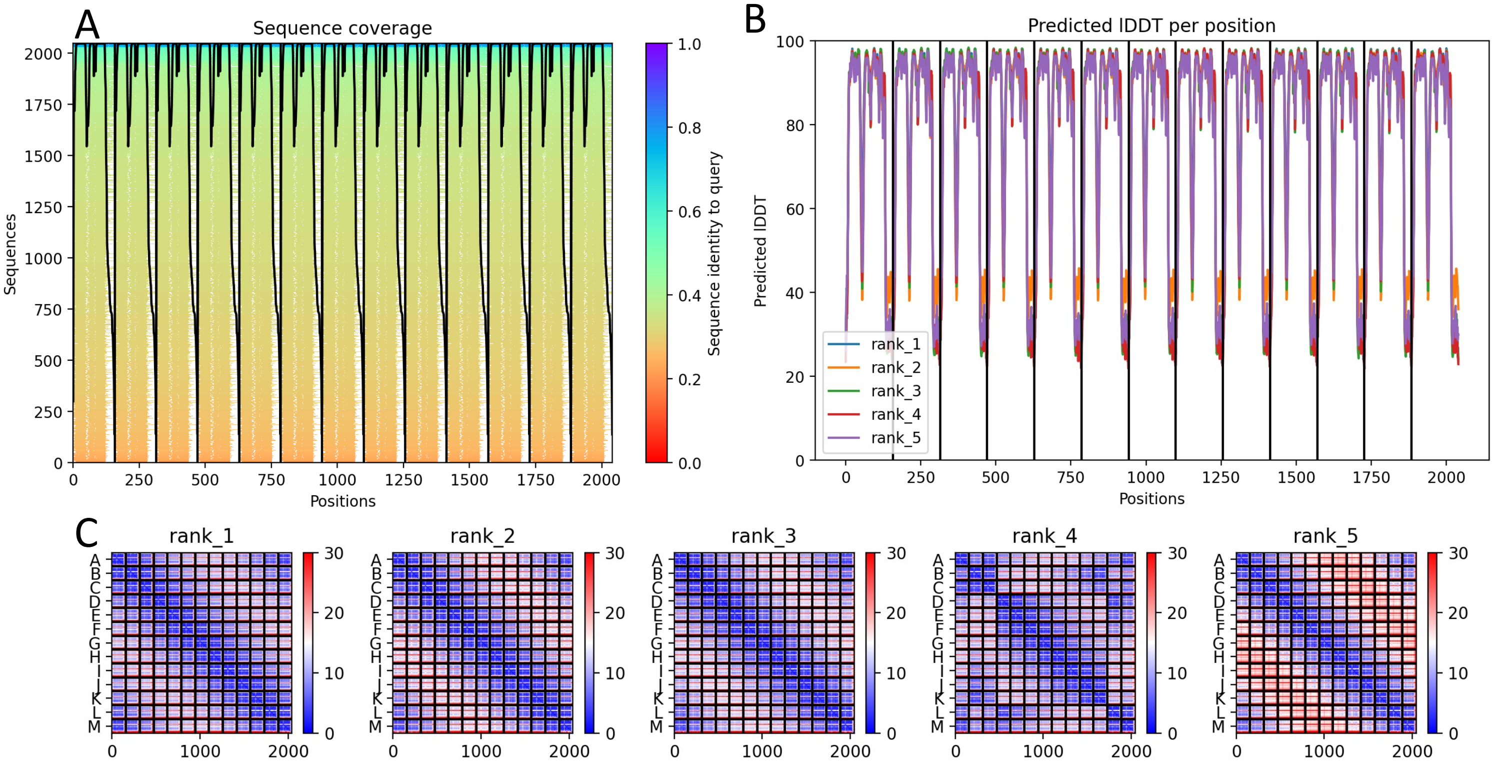


| | pLDDT | ptm | ipTM |
|---|---|---|---|
| model1 | 78.1 | 0.767 | 0.758 |
| model2 | 78.2 | 0.762 | 0.751 |
| model3 | 77.6 | 0.759 | 0.748 |
| model4 | 77.4 | 0.757 | 0.747 |
| model5 | 77.2 | 0.689 | 0.674 |

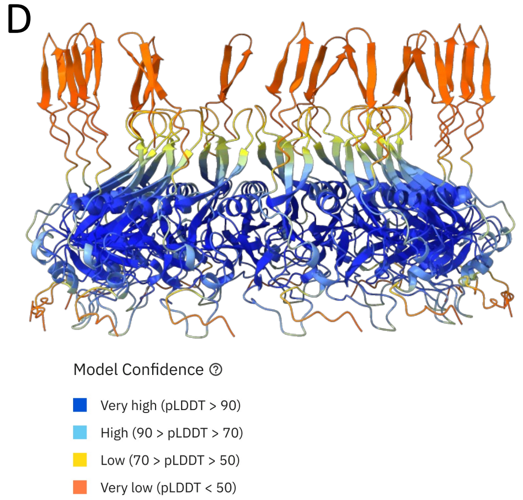


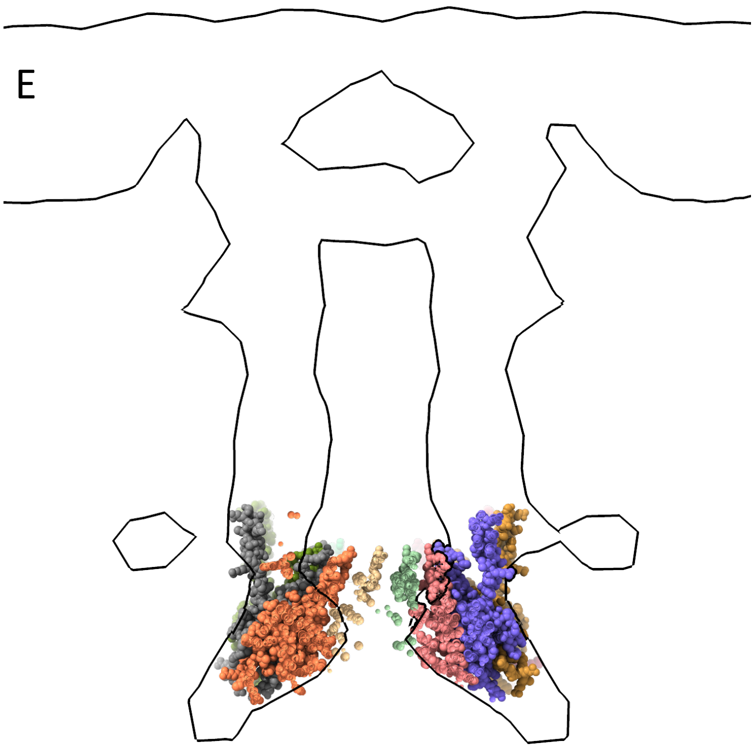


**Fig S9. | CpaB$_{141-297}$ C13 AlphaFold model.** (A) Sequence coverage, (B) pLDDT, (C) PAE, and table with metrics of the five different models for the C13 CpaB$_{141-297}$ multimer. (D) Best model colored by confidence. (E) Fitting of the CpaB C13 model, colored by chain identity, in the lower periplasmic ring of the non-piliated Δ*pilA* STA.

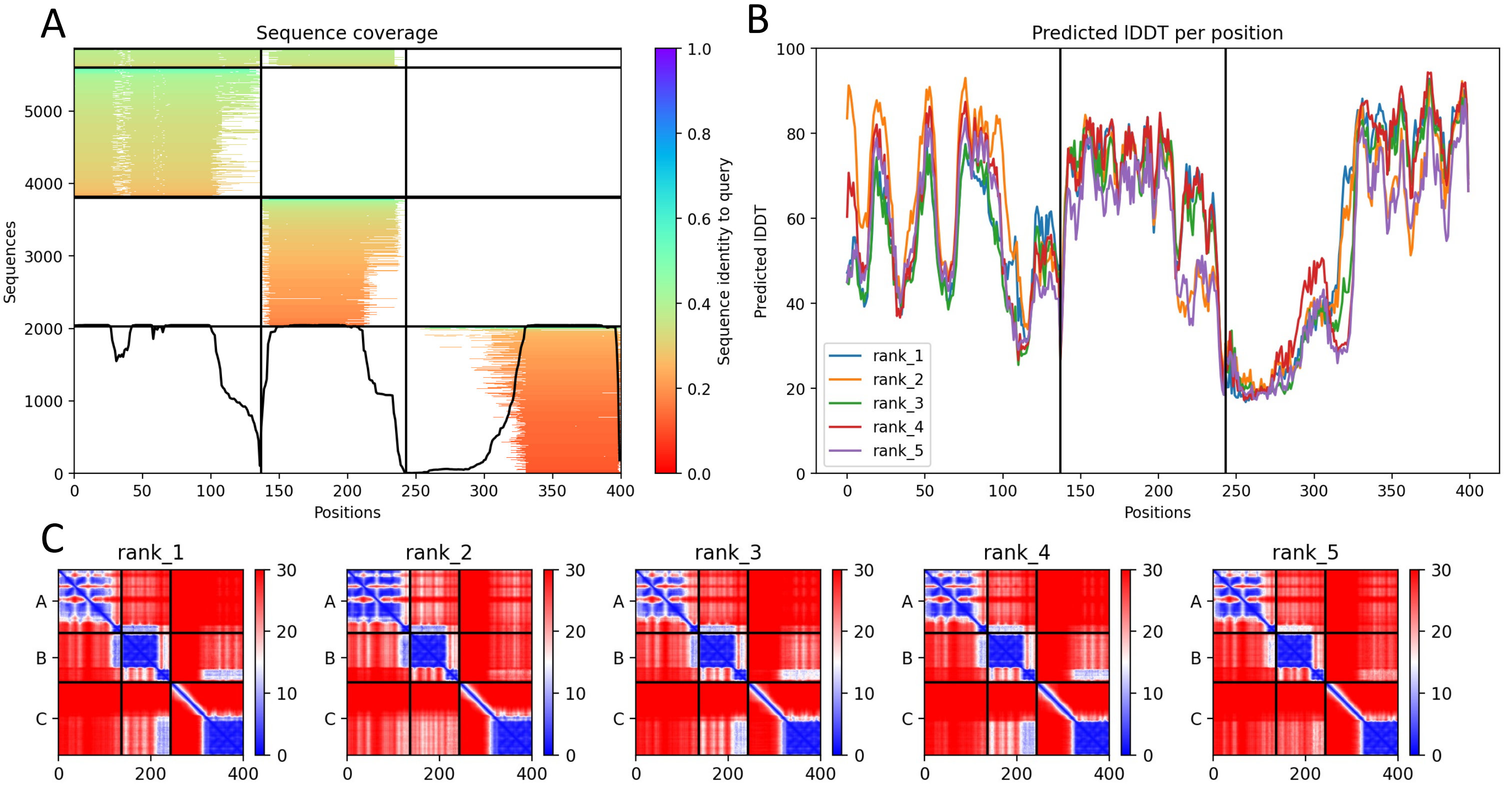


| | pLDDT | ptm | ipTM |
|---|---|---|---|
| model1 | 58.6 | 0.326 | 0.295 |
| model2 | 59.5 | 0.337 | 0.28 |
| model3 | 56.3 | 0.32 | 0.282 |
| model4 | 59.5 | 0.32 | 0.265 |
| model5 | 53.3 | 0.293 | 0.226 |

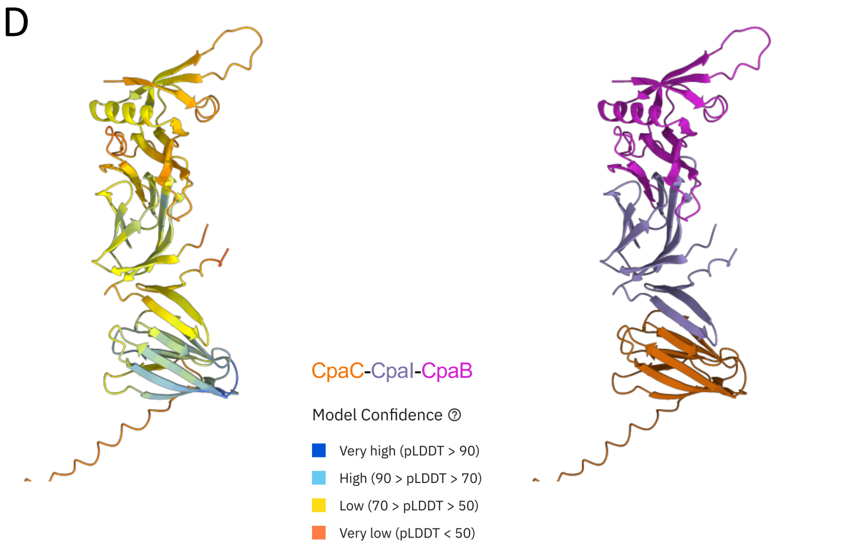


**Fig S10. | CpaC$_{N\text{-}term}$-CpaI-CpaB$_{141\text{-}297}$ AlphaFold Model.** (A) Sequence coverage, (B) pLDDT, (C) PAE, and table with metrics of the five different models for the CpaC$_{N\text{-}term}$-CpaI-CpaB$_{141\text{-}297}$ heterotrimer. (D) Best model colored by confidence and by chain identity.

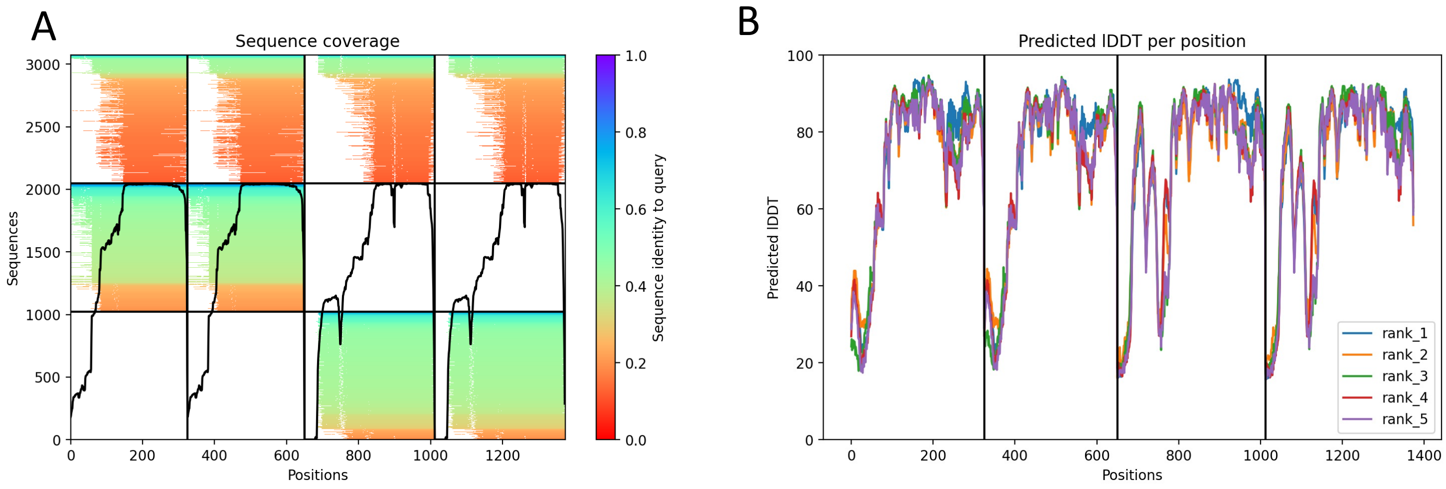


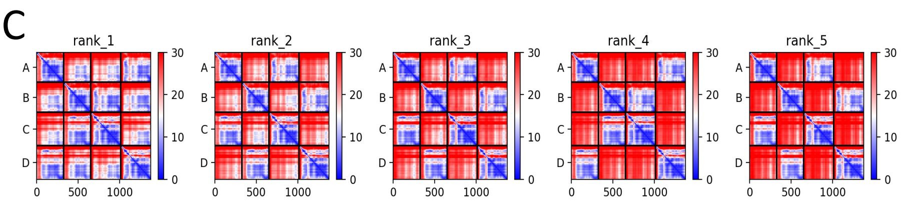


| | pLDDT | ptm | ipTM |
|---|---|---|---|
| model1 | 72.4 | 0.653 | 0.63 |
| model2 | 71.4 | 0.565 | 0.5 |
| model3 | 71.8 | 0.524 | 0.449 |
| model4 | 71.1 | 0.496 | 408 |
| model5 | 70.4 | 0.472 | 0.386 |

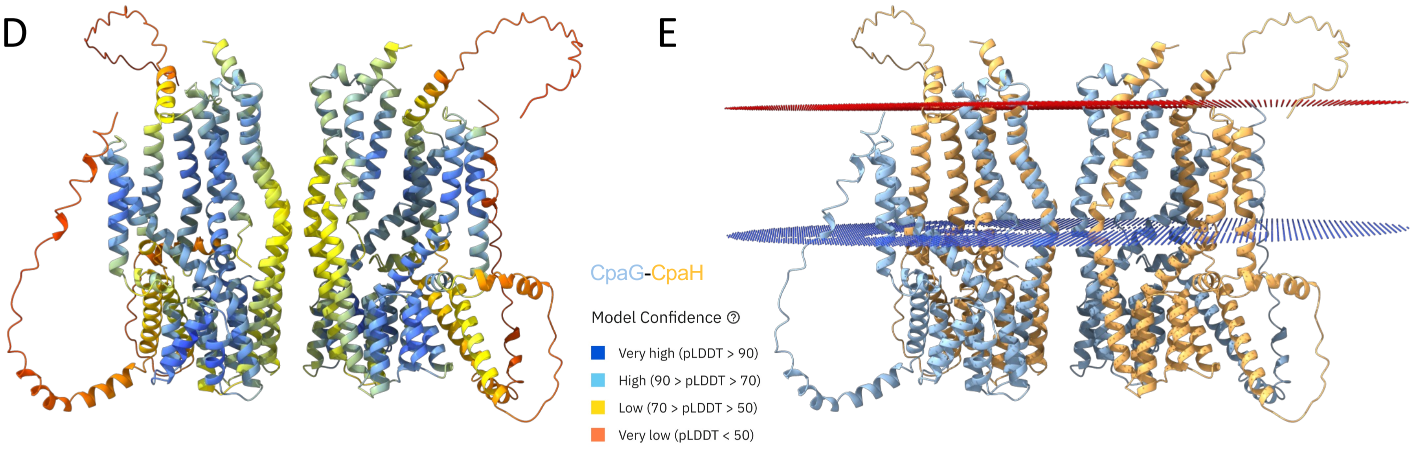


**Fig S11. | CpaG-CpaH AlphaFold model.** (A) Sequence coverage, (B) pLDDT, (C) PAE, and table with metrics of the five different models for the CpaG-CpaH heterodimer. (D) Best model colored by confidence and (E) by chain identity after prediction of the cytoplasmic membrane insertion (Cytoplasmic leaflet in blue, periplasmic leaflet in red).

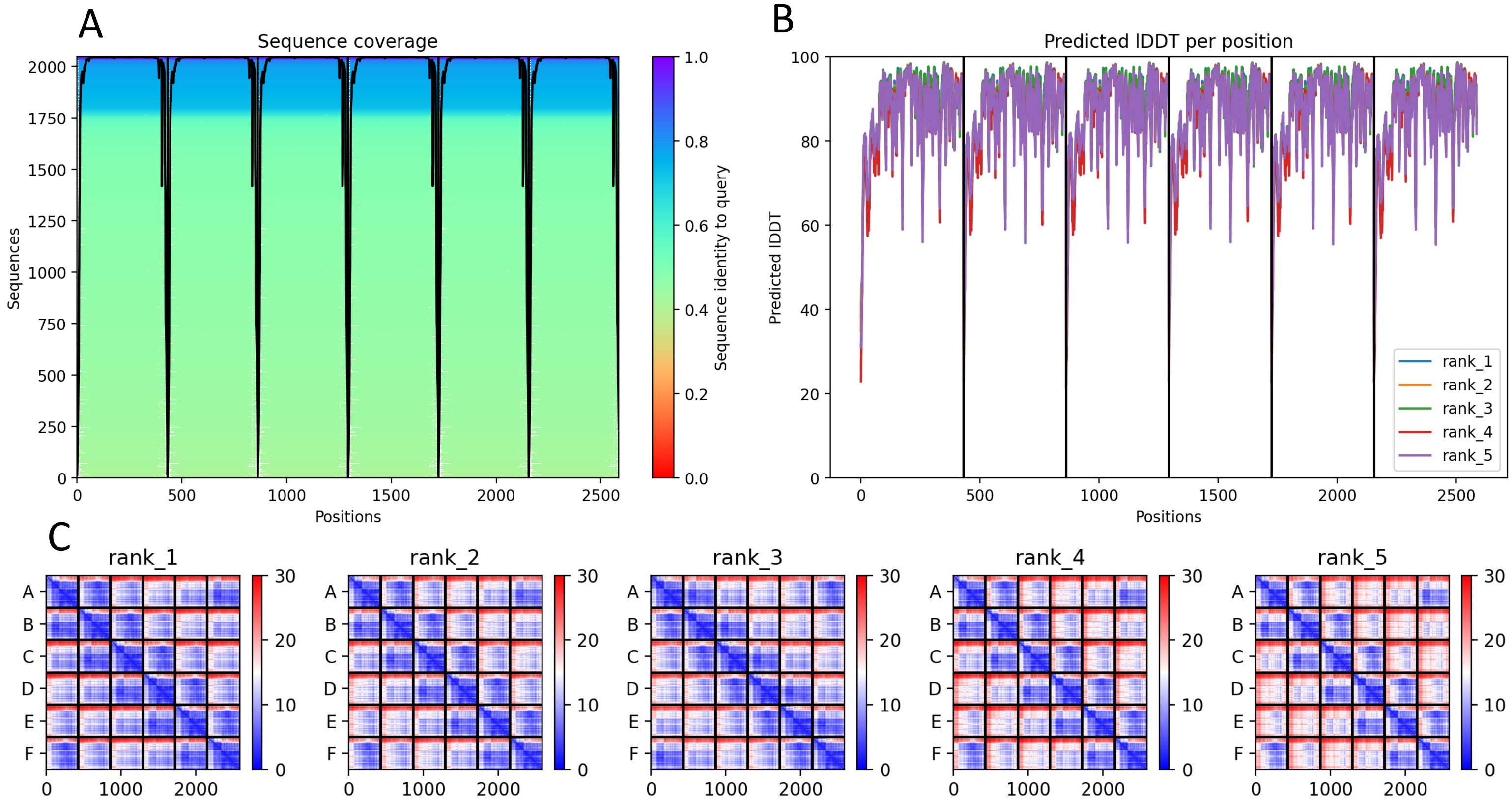


| | pLDDT | pTM | ipTM |
|---|---|---|---|
| model1 | 87.8 | 0.837 | 0.82 |
| model2 | 87.1 | 0.81 | 0.789 |
| model3 | 87.2 | 0.801 | 0.783 |
| model4 | 85.6 | 0.761 | 0.734 |
| model5 | 86.1 | 0.723 | 0.692 |

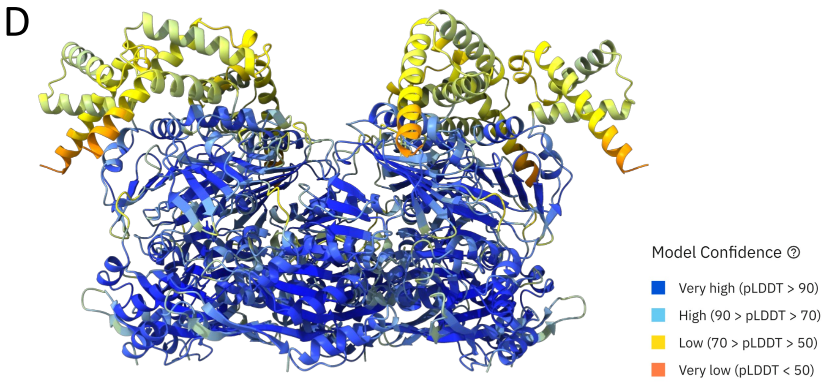


**Fig S12. | CpaF AlphaFold model.** (A) Sequence coverage, (B) pLDDT, (C) PAE, and table with metrics of the five different models for the CpaF hexamer. (D) Best model colored by confidence.

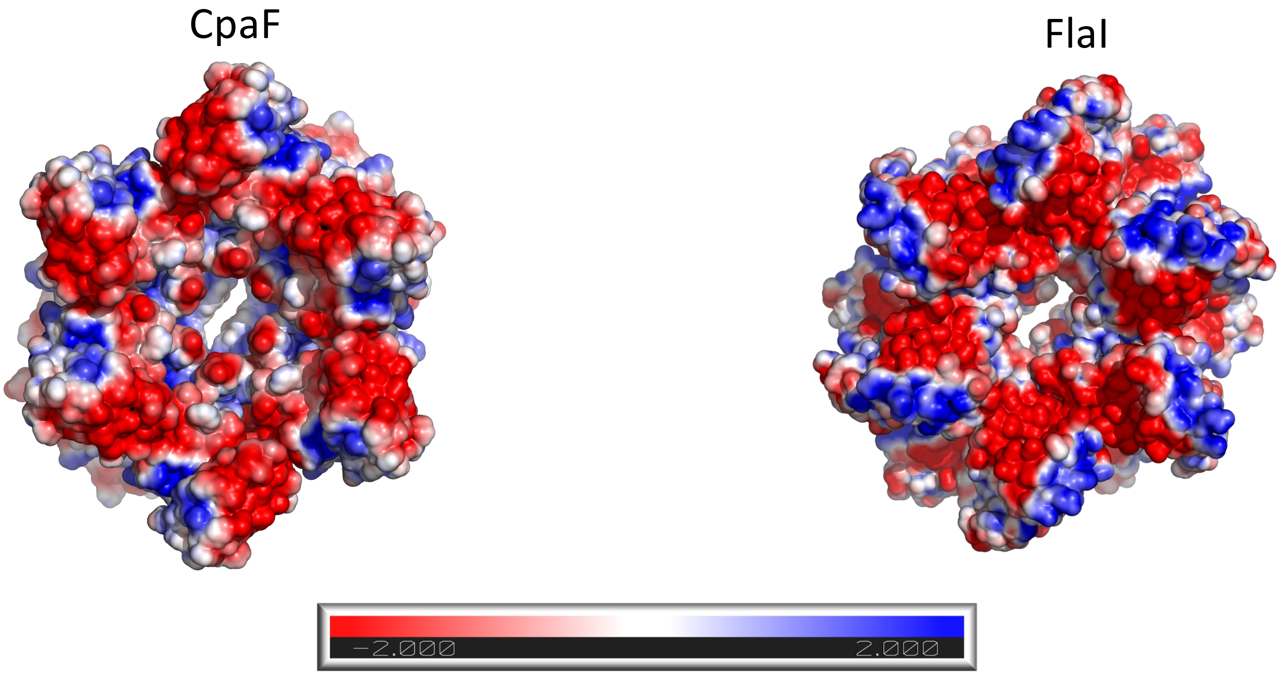


**Fig S13. | Electrostatic potential of *Cv*CpaF model and *Sa*FlaI.** Top view of the electrostatic potential CpaF model and FlaI (PDB: 4ihq) hexamers. The negative charge of this region is conserved between models and, in FlaI, interacts with the inner membrane platform FlaJ.

**Table S1**

| | |
|---|---|
| >caulo_**CpaG** | |
| FAFAGGDSASSKAAKRAQIIAKGGERQANVRAKAAANTPDARRQAILKALKDQDRKQKKASLSIAAR | 11.3 |
| >Asticcacaulis sp. AC466 WP_023457294.1 | |
| YAITGGGSGSSAALTKRTQTIGLGTSTRAKHRQKAQAKTPEERRKQISEQLKEAEKRERKARLTLRAR | 11.47 |
| >Brevundimonas naejangsanensis WP_368933249.1 | |
| WVLVGGDDSSGQAVKRAKVLATSRTESVAAAKRAAVANTPEARRKQIMVQLREVDRRERKARMTTSAK | 11.45 |
| >Woodsholea WP_019959982.1 | |
| FALIPSGNKRQTERVTQAVGAPLHRGRRHTAALDMAAQKKKQIQDSIKDIEERQKLARKRKVSLKAR | 11.5 |
| >Phenylobacterium zucineum WP_257838997.1 | |
| LPSTGRRRLARRARALGAGRKSAPEGRAVRLARPGATRLDALAQRLLPKPAVLRSR | 12.57 |
| >Parvularcula bermudensis WP_013300126.1 | |
| FALLGFDGGGATNKRVQSLSGNAKQAKRKTSAKEQTRDRRKTISDSLKKIEDKQKEANQKKKVSLDK | 10.38 |
| >Kordiimonas gwangyangensis WP_020398027.1 | |
| VIVLSNALSKGRKETRQKLERFKARFEAKSSLKGEGARSIRVNQQQSGLVASLSELLPRRDQIKRR | 11.66 |
| >Kiloniella laminariae WP_020593589.1 | |
| FILNPSSDISKTRLERVIHKAADGRSTAASNISIRRNMGGSGNPTLDKIVQQLTPRPELLRAK | 11.31 |
| >Parvibaculum lavamentivorans WP_011995151.1 | |
| FLFVGGESRSQKRVARVTRAGNARLVAVDGEVDNSEKRRKQMQETLKGLEEKQQEHKKRVSLNTR | 10.62 |
| >Sulfitobacter mediterraneus WP_025046443.1 | |
| LVAFGKSISLNSRVNRRLEMLEKGDRREEVLEKLRKEMQQHMKSKGIPLYSLLSEK | 9.99 |
| >Novosphingobium WP_271184746.1 | |
| WALLSGPSPKKESTRRLKSVRFRHSSNASDRLEAQMRRAVAARKPRIHQIAGSASRAEALALR | 12.08 |
| >Caenispirillum salinarum WP_009541532.1 | |
| MAFGGGTARKAVNRRLRLRASGKSSEEVMETLRRETPGQAAQGGPLGWLGR | 11.79 |

**Table S2**

| | |
|---|---|
| >caulo_**CpaH** | |
| PVMRDNNLEGRLKSVANRREELRRRSRQSISTRAPGTAGGTLRHQDEGLYKNVVERLQLSRLLEDPKVVEKLAQAGFRGPKPVSTF | 11.1 |
| >Asticcacaulis sp. AC466 (uniprot:V4NFP1) | |
| SMTSGEQLDKRLKAVAQRREELRRQSRQQIVKEQSGGLRHKDDGMARGFVDKFNLRKLLEDPKVAENLAMAGFRGPKPISTF | 10.41 |
| >Brevundimonas naejangsanensis WP_003168046.1 | |
| SFVGGGPRLDKRMKAVAERREELRRRSRQALRGGGGETGGGLRRTDDSFRSRVVARLNLVKLLEDPKVAENMTQAGFRGPRPLNTF | 11.67 |
| >woodsho WP_019959981.1 | |
| PLMGGDSLDKRIKAVHVRREELRRKSREAIDNAHKPQNQRLRGDQGKGLYRNVVEKLNLQKLLEDPDLKDKLVQAGLRGQGPIFTF | 10.15 |
| >Phenylobacterium zucineum WP_257838998.1 | |
| ALSPGDPVAVRARAHARRRQELRAELLRPRRSERARSVGLARRLVEALNLNRGDEAKKAASLLVQAGLRTSDATHVF | 11.83 |
| >Parvularcula bermudensis WP_013300127.1 | |
| PLFADNGLDKRVKKVADYKERLRQQSRAELAGSRKPGSLRNATAGGMGKRLVEQLKLFEIFDAAAARKQLMMAGLRGERPVFTY | 10.52 |
| >Kordiimonas gwangyangensis WP_020398028.1 | |
| AALVRDPMKGRVKSLQGRRDALKAGLMTTSKRRSPVKKVDSVGALRALADKFQLLQKEQTAKVSQKLTQAGLRSSDAVVIY | 11.08 |
| >Kiloniella laminariae WP_020593590.1 | |
| SLIEKDPSSKKAKELSELRKAMRGDYLTPTQRKRGNENSSTYMKKVVEFFKLLKNEQASKYAFHLSQAGRRTPEALYRH | 10 |
| >Parvibaculum lavamentivorans WP_011995152.1 | |
| PLLNTDKLSTRMKYVSTEREAMRARAREALAQEKTRLRHTPKGFIKQVVEKFASGNMLENAAVKEKLRQAGFRGPGPMYTF | 10.53 |
| >Sulfitobacter mediterraneus WP_025046444.1 | |
| MVNQPEDPLDKLKRDRTAPRKGDGQIATLRQKEGNAQLQKYATFLEPKDEGELSQKQLELRQAGYQSRDAVRMF | 9.1 |
| >Novosphingobium WP_271184745.1 | |
| AITIRDPMAKRVKALNERRDALKAGLVTTARKRQSLVRRNDTTDRMGAFLGGAKMLQDSQIKVIQQKLAQAGIRNKEWAVAV | 11.67 |
| >Caenispirillum salinarum WP_009541533.1 | |
| AVYGIAGLRSGVAARRFQRGGGRGTGGAAAAPDNDVSLRPDGAFQKVEAFLQPYGETLEPQKVEERKRIRLTLLRAGFPQANAITIY | 10.25 |
